# Strand asymmetry in the repair of replication dependent double-strand breaks

**DOI:** 10.1101/2024.06.17.598707

**Authors:** Michael T Kimble, Aakanksha Sane, Robert JD Reid, Matthew J Johnson, Rodney Rothstein, Lorraine S Symington

## Abstract

Single-strand breaks (SSBs) are one of the most common endogenous lesions and have the potential to give rise to cytotoxic double-strand breaks (DSBs) during DNA replication. To investigate the mechanism of replication fork collapse at SSBs and subsequent repair, we employed Cas9 nickase (nCas9) to generate site and strand-specific nicks in the budding yeast genome. We show that nCas9-induced nicks are converted to mostly double-ended DSBs during S-phase. We find that repair of replication-dependent DSBs requires homologous recombination (HR) and is independent of canonical non-homologous end joining. Consistent with a strong bias to repair these lesions using a sister chromatid template, we observe minimal induction of inter-chromosomal HR by nCas9. Using nCas9 and a gRNA to nick either the leading or lagging strand template, we carried out a genome-wide screen to identify factors necessary for the repair of replication-dependent DSBs. All the core HR genes were recovered in the screen with both gRNAs, but we recovered components of the replication-coupled nucleosome assembly (RCNA) pathway with only the gRNA targeting the leading strand template. By use of additional gRNAs, we find that the RCNA pathway is especially important to repair a leading strand fork collapse.

## INTRODUCTION

Cells must faithfully replicate the genome and repair DNA damage in each cell cycle to maintain genome integrity. It is estimated that tens of thousands of single-strand breaks (SSBs) occur each day in human cells, making them one of the most common endogenous lesions.^1,2^ These lesions can arise from oxidative damage and as transient intermediates in DNA metabolic processes, such as topoisomerase I cleavage or base excision repair. SSBs have the potential to give rise to DSBs during replication by a mechanism that involves fork collapse. Unidirectional fork collapse at an SSB is proposed to result in a single-ended double strand break (seDSB) that is subsequently repaired by homologous recombination (HR) with the intact sister chromatid.^3–5^ Thus, investigating how SSBs are repaired has the potential to yield insight into the mechanisms of sister-chromatid recombination, which cannot be easily studied with endonucleases that cleave both sister chromatids. In particular, it is unknown if the genetic requirements to repair replication-dependent DSBs are the same as for DSBs induced outside of S-phase.

Camptothecin (CPT), a commonly used genotoxic agent, creates stabilized protein-bound SSBs by trapping Top1 cleavage complexes, and therefore potentiates replication-associated DSBs.^6^ The repair pathway usage of CPT-induced damage closely matches that of spontaneous lesions, supporting the physiological relevance of this type of DNA lesion.^7^ To study this type of DNA damage in a site-specific manner, others have developed the Flp-nick system to introduce a single protein-bound nick in the *Saccharomyces cerevisiae* genome that is converted to a seDSB during S phase.^8^ Cell survival in response to Flp-mediated nicking is reduced in the absence of HR factors,^4,8,9^ consistent with the need for sister chromatid strand invasion to complete DNA replication. In a recent study, the Flp-nick system was integrated into the genome of mouse cells and shown to stimulate RAD51-dependent recombination between closely linked repeats.^10^ A minimal HO endonuclease recognition site that is mostly nicked by HO has been used to study the repair of replication-associated DSBs in a plasmid context.^11^ In the plasmid system, the nick in converted to a double-ended DSB (deDSB) during replication; however, it is unclear whether this is due to fork convergence within the small replicon or to bypass of the lesion by the replicative helicase (CMG).^12^ While these systems offer the ability to study a unique replication-associated break, efficiency of cleavage is relatively low, both require introduction of an enzyme recognition site into the genome, and they lack strand specificity. By contrast, Cas9 nickase (nCas9) offers the ability to induce nicks in a site and strand-specific manner in the native genome with the versatile design of a single gRNA.

There are two versions of nCas9, depending on which nuclease site remains active. The Cas9^D10A^ variant contains a point mutation in the RuvC domain and cleaves the gRNA target strand using the HNH domain, while the HNH domain of Cas9^H840A^ is inactivated and it cleaves the non-target strand via the RuvC domain.^13^ Cas9 nickase has been used in genome editing, proving particularly useful in the paired nick strategy and prime editing.^14,15^ More recently, it has also been employed to study the recombinogenic capacity of replication-associated breaks in mammalian cells. ^16–18^ These studies have shown that both nickases stimulate intrachromosomal HR between repeat sequences with Cas9^D10A^ showing highest activity. ^16,17^ The ratio of long-tract to short-tract gene conversion (LT/STGC) products in response to nCas9 expression is higher than observed using Cas9 or I-SceI, consistent with more extensive DNA synthesis associated with HR in S-phase or to break-induced replication (BIR) initiated at a seDSB.^16,19^

Here we establish use of Cas9 nickase in *Saccharomyces cerevisiae* to study repair of collapsed replication forks. We show that nick induction with Cas9^D10A^ leads to replication-dependent DSBs. However, contrary to expectation, these are mostly deDSBs instead of seDSBs. Based on the genetic requirements for cell survival, we demonstrate that replication-dependent breaks are repaired through HR, while DSBs generated by Cas9 rely on non-homologous end joining (NHEJ). By systematic screening of the yeast non-essential gene deletion library, we identified the replication-coupled nucleosome assembly (RCNA) pathway as essential for repair of collapsed replication forks. Previous screens in both yeast and humans have identified mutants of the RCNA pathway by their sensitivity to CPT, methyl methanesulfonate (MMS) and to unscheduled RNaseH2 expression, thereby implicating this pathway in maintaining genome integrity during replication stress.^20–23^ Our studies reveal that the RCNA pathway is especially important to facilitate repair of breaks that form on the leading strand template, whereas a nick on the lagging strand template is tolerated.

## RESULTS

### Cas9 nickase creates replication-dependent DSBs

To study the repair of broken replication forks, we established yeast strains expressing nCas9 with a gRNA designed to target a unique sequence in the genome. We generated two versions of nCas9, Cas9^D10A^ and Cas9^H840A^, each with a single amino acid change to ablate one of the two Cas9 nuclease active sites.^13^ In our system, Cas9 or nCas9 is conditionally expressed by addition of *β*-estradiol to the growth medium^24,25^ and the gRNA is constitutively expressed from a Tyr-tRNA promoter. We designed gRNAs to target sequences near the efficient, early firing *ARS607* origin on chromosome VI (Figure 1A and S1A),^26,27^ and tested sensitivity to induction of Cas9^D10A^ by plating efficiency of cells on medium with or without *β*-estradiol. We compared growth inhibition by Cas9^D10A^ in a wild-type (WT) and an *mre11Δ* background with gRNAs targeting either centromeric proximal (gRNAs 1-3) or telomeric proximal (gRNAs 6-10) to *ARS607* (Figure S1B). A previous study had shown that induction of the Flp-nick system inhibits growth of Mre11 or Rad51-deficient cells;^8^ therefore, we reasoned that efficient gRNAs would sensitize HR-deficient cells to nCas9 expression. This analysis identified gRNA1 and gRNA6 as highly efficient in growth inhibition of *mre11Δ* cells (Figure S1B). Based on origin timing and proximity,^26,27^ gRNA1 is predicted to direct Cas9^D10A^ to the lagging strand template, whereas gRNA6 should cause a leading strand fork collapse. Further screening identified gRNA14 and gRNA20 to efficiently cleave on the opposite strands to gRNA1 and gRNA6, respectively (Figure S1C). To verify DNA cleavage by the gRNAs selected, each was expressed with Cas9 in asynchronous cultures and genomic DNA isolated for Southern blot analysis. In each case, ∼50% of the target site was cut after a 90-min Cas9 induction (Figure S1D).

**Figure 1.**
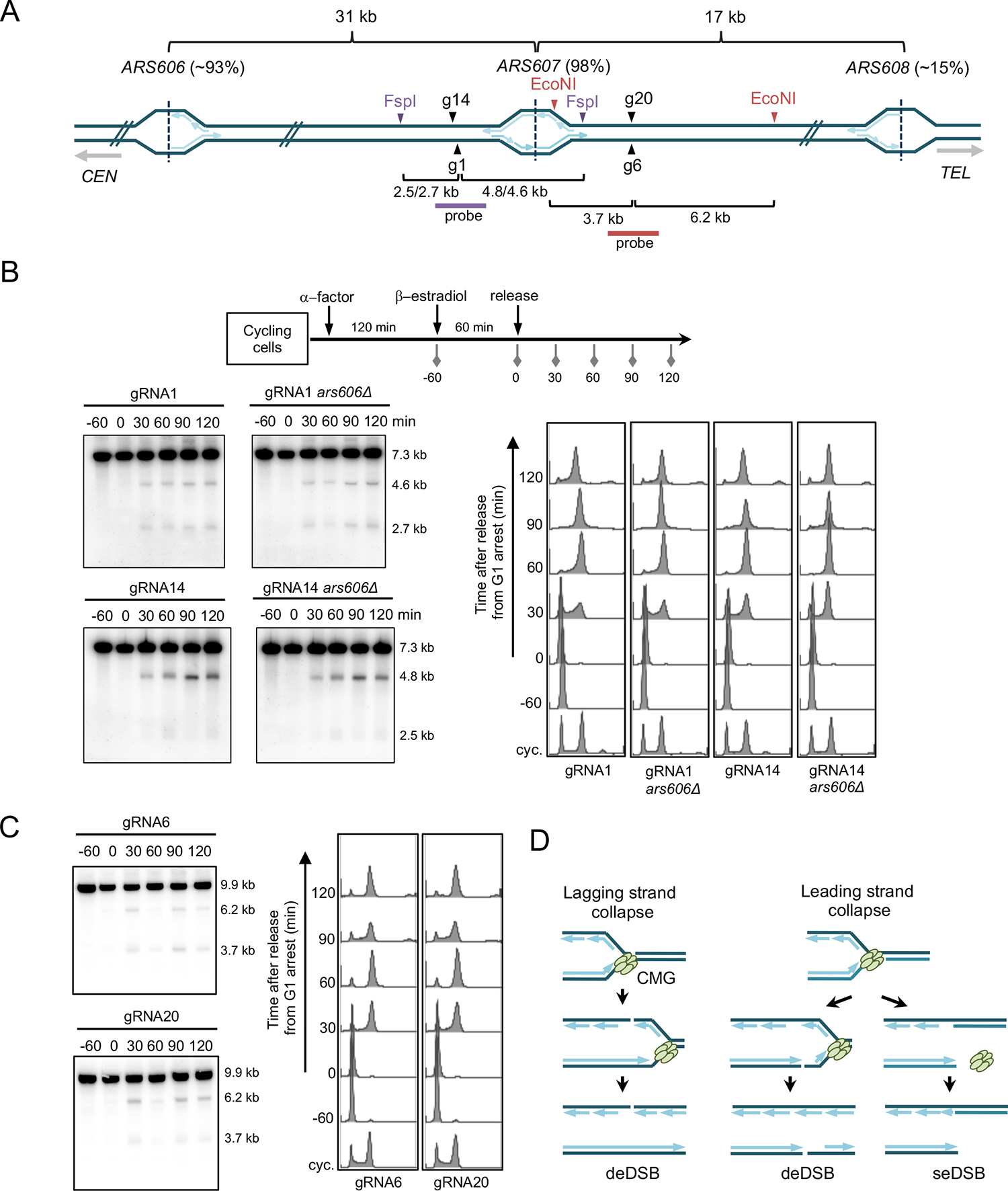
Cas9^D10A^ induces replication-dependent DSBs. A. Schematic of the right arm of chromosome VI indicating the locations of confirmed replication origins (*ARS* elements) and the gRNA (g) cleavage sites. The activity of each *ARS* in the W303 strain background ^27^ is shown in parentheses. The expected DNA fragment sizes from cleavage with Cas9 or fork collapse at Cas9^D10A^-induced nicks, followed by digest with FspI or EcoNI restriction enzymes are shown. Note that the sequence recognized by g14 is 200 bp centromeric to g1. The location of hybridization probes spanning the gRNA sites are indicated by the purple and dark red lines. B. Schematic for nCas9 induction in synchronized cells and Southern blots of neutral gels showing replication-dependent DSBs with g1 and g14, along with FACS profiles from the same samples. C. Southern blots of neutral gels showing replication-dependent DSBs with g6 and g20, along with FACS profiles from the same samples. D. Model for replication fork collapse at nicks induced by Cas9^D10A^ on the leading or lagging strand template.

Based on data from nicks induced by CPT and Flp-nick, nCas9 should yield a seDSB when the replication fork collapses at the nick.^8,28,29^ We synchronized *MAT***a** cells in the G1 phase with *α*-factor (t-60), induced expression of Cas9^D10A^ with gRNA1 or gRNA14 for 60 min, released cells from the G1 arrest and isolated genomic DNA before release (t 0) and at 30 min intervals after release into S phase for Southern blot analysis (Figure 1B). We expected that a replication fork initiated from *ARS607* would generate a 4.6 or 4.8 kb band for gRNA1 or gRNA14, respectively, reflecting a fork moving right to left resulting in collapse (Figure 1A). DSBs were detected 30 min after release into S-phase consistent with replication dependence (Figure 1B), since no breaks were detected in G1-arrested cells 60 min after Cas9^D10A^ expression (Figure 1B, t 0). Although we observed the expected seDSB with gRNA14 (leading strand break), a double-ended DSB (deDSB) was generated at the lagging strand nick produced by gRNA1. We considered the possibility that delayed repair of a lagging strand fork collapse might allow time for a converging fork to arrive, converting the seDSB to deDSB. To test this idea, we deleted *ARS606*, an efficient origin located 31 kb centromere proximal to *ARS607*; however, the deDSB persisted in the *ars606Δ* strain expressing Cas9^D10A^ with gRNA1. The next origin distal to *ARS606* is located on the left arm of chromosome VI and would not be expected to reach the nick site within 30 min after release of cells from the G1 arrest.^26,27^ This result suggests that the replicative helicase continues translocation on the leading strand template after encountering the lagging strand nick, generating what would appear as a deDSB by continued lagging strand synthesis (Figure 1D). Of note, one end of the deDSB is predicted to be blunt and the other to have an overhang of up to the size of an Okazaki fragment.^5^

To determine the generality of these findings, we performed a similar genomic DNA analysis for synchronized cells expressing Cas9^D10A^ and gRNA6 or gRNA20 (Figure 1A, C). These guides target sequences 4 kb telomeric to *ARS607* and 13 kb from *ARS608*, which is reported to be an inefficient origin.^26,27^ In agreement with the data obtained with gRNA1 and gRNA14, DSBs were detected 30 min after release from G1 indicating replication dependence. However, we observed deDSBs for both leading and lagging strand nicks. To determine whether the deDSB signal results from a converging fork, we employed a strain deleted for two annotated origins, *ARS608* and *ARS609* (57 kb telomeric to *ARS607*) (Figure S2A).^30,31^ Surprisingly, deDSBs were still detected with both gRNAs (Figure S2B), raising the possibility that CMG can translocate over the nick on the leading strand template (Figure 1D). We note that gRNA6 is 60% G-rich and this could potentially increase the stability of the gRNA/DNA hybrid post cleavage to allow bypass by CMG and continued unwinding of the leading strand. Alternatively, replication fork collapse could activate dormant origins as has been shown for HO-endonuclease-induced DSBs during S-_phase._32

### Homologous recombination is essential for repair of collapsed replication forks

The physical analysis indicates that Cas9^D10A^ can generate replication-dependent single-end or deDSBs. Repair of both types of breaks is expected to require end resection and Rad51-mediated invasion of the sister chromatid. However, a deDSB would have greater potential for repair by NHEJ and different modes of D-loop resolution compared with a seDSB. We tested sensitivity of WT cells, or mutants defective for NHEJ (*dnl4Δ*), end resection (*mre11Δ*), or strand invasion (*rad51Δ*) to induction of nCas9 or Cas9 by plating efficiency of cells on medium with or without *β*-estradiol. Cas9^D10A^ induction with gRNA1 or gRNA6 did not impair growth of haploid WT or *dnl4Δ* cells, confirming that the DSBs detected by physical assays do not arise in G1 cells (Figure 2A). However, viability of *mre11Δ* and *rad51Δ* cells was reduced by ∼1000-fold indicating that repair is highly dependent on HR. Cas9^H840A^ expression was slightly less toxic to HR-deficient strains compared to Cas9^D10A^, which is consistent with previous findings that Cas9^H840A^ is not as effective as Cas9^D10A^ in stimulating HR.^16,17^ Cas9^H840A^ could be a less efficient nickase or the nick is more readily repaired by a non-HR process. For this reason, we primarily used Cas9^D10A^ for subsequent analyses. Growth inhibition in *mre11Δ* and *rad51Δ* backgrounds requires a targeting gRNA, as a scrambled gRNA1 in combination with Cas9^D10A^ did not reduce cell viability. A catalytically inactive Cas9 (dCas9) in combination with gRNA1 or gRNA6 also did not confer sensitivity, confirming that the growth defect observed for HR-deficient strains is due to nicking and not simply Cas9 binding to DNA (Figure 2A). As expected, Cas9 expression with either gRNA1 or gRNA6 reduced viability of WT cells due to the inefficiency of error-prone NHEJ in yeast and was lethal in the absence of Dnl4. ^33^ In budding yeast, the Mre11-Rad50-Xrs2 (MRX) complex is essential for NHEJ and, consequently, growth of *mre11Δ* was inhibited by Cas9 expression with both gRNAs. Interestingly, Cas9 expression was less toxic than Cas9^D10A^ in the *rad51*Δ background. Although deDSBs are generated with gRNA1 and gRNA6 (Figure 1B), repair is still highly dependent on Rad51 suggesting that NHEJ repair of a replication-dependent DSB is inefficient. As noted above, one end of a replication-dependent deDSB is predicted to have a ssDNA overhang and to be less amenable to repair by canonical NHEJ. Together, these data show that a single broken replication fork is lethal in HR-deficient cells and that these breaks are repaired in a different way to Cas9-induced DSBs.

**Figure 2.**
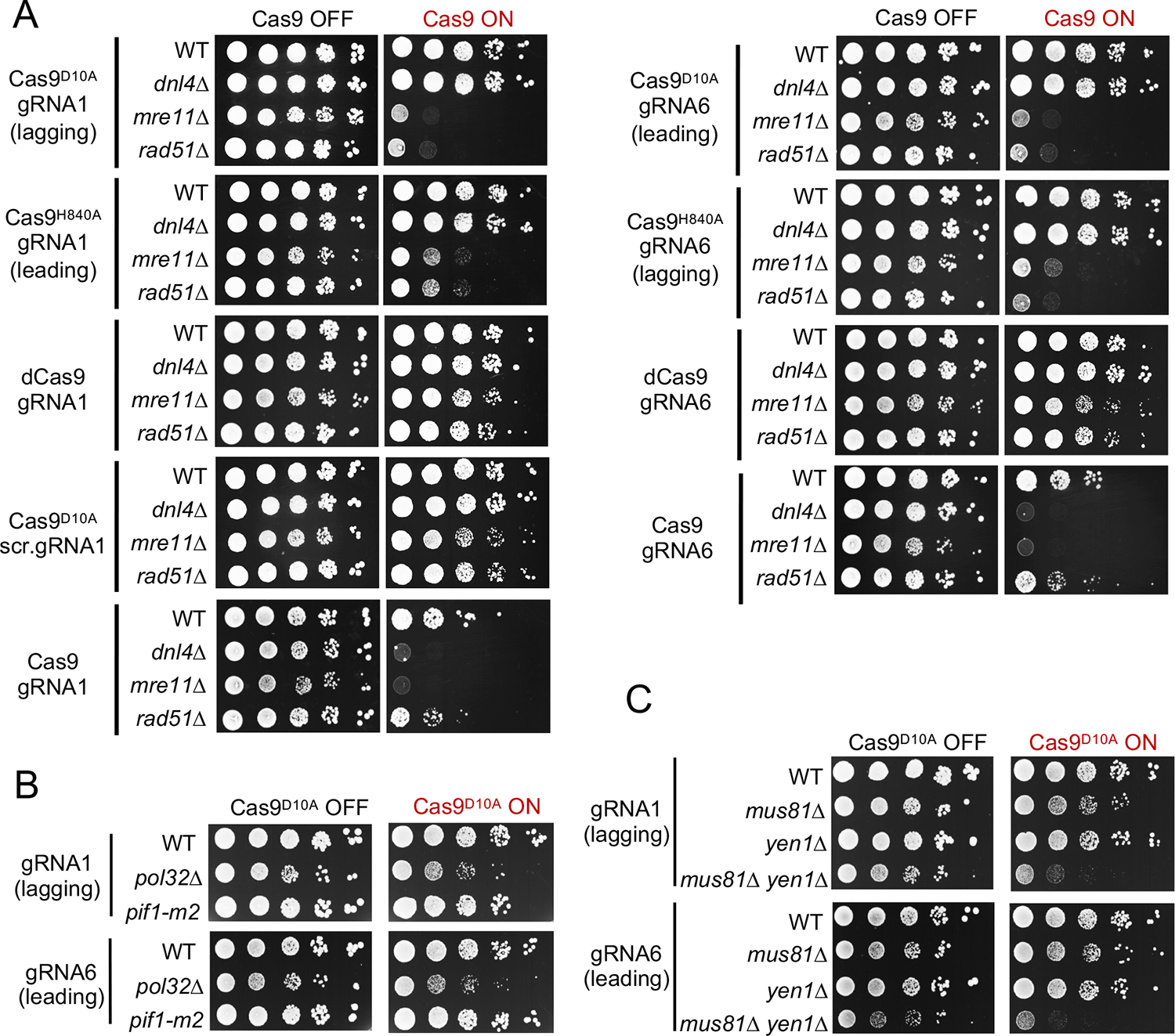
Homologous recombination is essential for repair of collapsed replication forks. A. Sensitivity of WT, *mre11Δ*, *rad51Δ* and *dnl4Δ* strains to expression of Cas9^D10A^, Cas9^H840A^, dCas9 or Cas9 paired with gRNA1 or gRNA6. Ten-fold serial dilutions of the indicated strains were spotted on medium +/- *β*-estradiol and grown for 3 days. B. Sensitivity of WT, *pol32Δ*, and *pif1-m2* strains to expression of Cas9^D10A^ paired with gRNA1 or gRNA6. Ten-fold serial dilutions of the indicated strains were spotted on medium +/- *β*-estradiol and grown for 3 days. C. Sensitivity of WT, *mus81Δ*, *yen1Δ* and *mus81Δ yen1Δ* strains to expression of Cas9^D10A^ paired with gRNA1 or gRNA6. Ten-fold serial dilutions of the indicated strains were spotted on medium +/- *β*-estradiol and grown for three days. Spot assays were performed for three independent transformants of each strain and representative plates are shown.

The MRX complex (MRN in human) promotes end resection, has a structural role in tethering DNA ends, and activates the Tel1/ATM kinase.^34^ To assess the contribution of end resection by MRX in collapsed fork repair, we tested growth inhibition of cells lacking resection nucleases to Cas9^D10A^-induced nicking. We found that nuclease-dead *mre11-H125N* cells were only mildly sensitive to expression of Cas9^D10A^, indicating that collapsed fork repair does not depend on Mre11’s nuclease activity (Figure S3A). This finding is consistent with previous studies in yeast showing that clean DSB ends can be processed by the other nucleases in the absence of Mre11 nuclease activity.^35^ Mre11 may also be required due to its role in recruiting the long-range resection proteins Exo1, Sgs1 and Dna2 to DSB ends ^36^. Therefore, we tested sensitivity to Cas9^D10A^ induction in *exo1Δ sgs1Δ* cells and observed a strong growth inhibition (Figure S3A). Deletion of either *EXO1* or *SGS1* alone did not cause a growth defect, consistent with redundancy between the long-range resection nucleases.^35^ The role of long-range resection is not immediately obvious, given that it dispensable for HR between closely linked repeats,^37^ and it has previously been reported that resection tracts at collapsed forks are relatively short.^9^ However, long-range resection may be required for efficient cohesin loading at a collapsed fork, since it has recently been shown that long-range resection is required for recruitment of the cohesin loader, Scc2, at DSBs.^38^

A study using Flp-nick to generate seDSBs showed that BIR is largely dispensable for repair.^4^ BIR requires the non-essential subunit of DNA polymerase *δ*, Pol32/POLD3, and to a lesser degree, the Pif1 helicase.^39,40^ Elimination of Pol32 or nuclear Pif1 (*pif1-m2*) had little effect on survival in the presence of a gRNA1 or gRNA6 nick, with a slightly stronger phenotype for *pol32Δ* (Figure 2B). Thus, BIR does not contribute significantly to repair of the Cas9^D10A^-induced replication fork collapse, consistent with generation of mainly deDSBs. A previous study showed that Holliday junction (HJ) resolution is important for repair of Flp nick-induced seDSBs.^4^ We found that growth of a *mus81Δ yen1Δ* mutant was severely impaired after Cas9^D10A^ expression with gRNA1 or gRNA6 (Figure 2C). The role of Mus81-Mms4 and/or Yen1 is likely to cleave the D-loop intermediate or HJ formed by second end capture at a deDSB. Therefore, repair of replication-associated DSBs does not require extensive BIR synthesis or Mre11 nuclease activity, but does require the MRX complex, long-range end resection, and HJ resolvases.

### Cas9 nickase stimulates sister chromatid recombination

Previous work in mammalian cells showed that nCas9 is less effective in stimulating HR between tandem repeats of GFP than Cas9 or I-SceI.^16,17,41^ To determine whether this is the case in yeast, we employed an *ade2* direct repeat assay ^42^ to compare recombination frequencies in response to Cas9 or nCas9 expression. A gRNA was designed to target the I-SceI cut site within the *ade2-I* allele (Figure 3A). There is a confirmed origin near the *ADE2* promoter that is duplicated within the recombination reporter; thus, replication is predicted to occur from right to left through the *ade2-I* allele. The gRNA should direct Cas9^D10A^ cleavage to the lagging strand, while Cas9^H840A^ should nick the leading strand template. Log-phase cells were plated on medium with or without *β*-estradiol and the Ade^+/-^ phenotype assessed by the colony color of surviving colonies (Ade^+^ colonies are white whereas Ade^-^ colonies are red). Retention of the *TRP1* marker located between the repeats was determined by replica plating to medium lacking tryptophan. A nCas9-induced deDSB could result in an Ade^+^ event through short tract gene conversion (STGC) with the *ade2-n* allele (intrachromatid or unequal sister chromatid interaction), whereas equal sister chromatid recombination would restore the *ade2-I* allele, potentiating a second round of nicking and fork collapse. Repair using the *ade2-n* allele would eliminate the gRNA binding site, preventing iterative nicking. A seDSB that invaded the *ade2-n* allele and initiated BIR would be predicted to copy the *ade2-n* allele generating Ade^-^ recombinants, since the *ade2-I* mutation is close to the 5*′* end of the ORF and the *ade2-n* mutation is near the 3*′* end. Ade^+^ recombinants could be generated by a template switch mechanism, but these events are expected to be rare ^43,44^. After expression of Cas9^D10A^ or Cas9^H840A^ and I-SceI gRNA we found that ∼100% of cells survived and most of the survivors were Ade^+^ Trp^+^, consistent with repair of a deDSB by short tract gene conversion (Figure 3B). Induction of a DSB with Cas9 resulted in a modest survival defect and a majority of events were Ade^+^ Trp^+^ (Figure 3B), similar to products formed by nCas9. Thus, nCas9 is proficient for induction of recombination between tandem repeats, and the predominance of STGC products is consistent with repair of a deDSB by conservative synthesis-dependent strand annealing.

**Figure 3.**
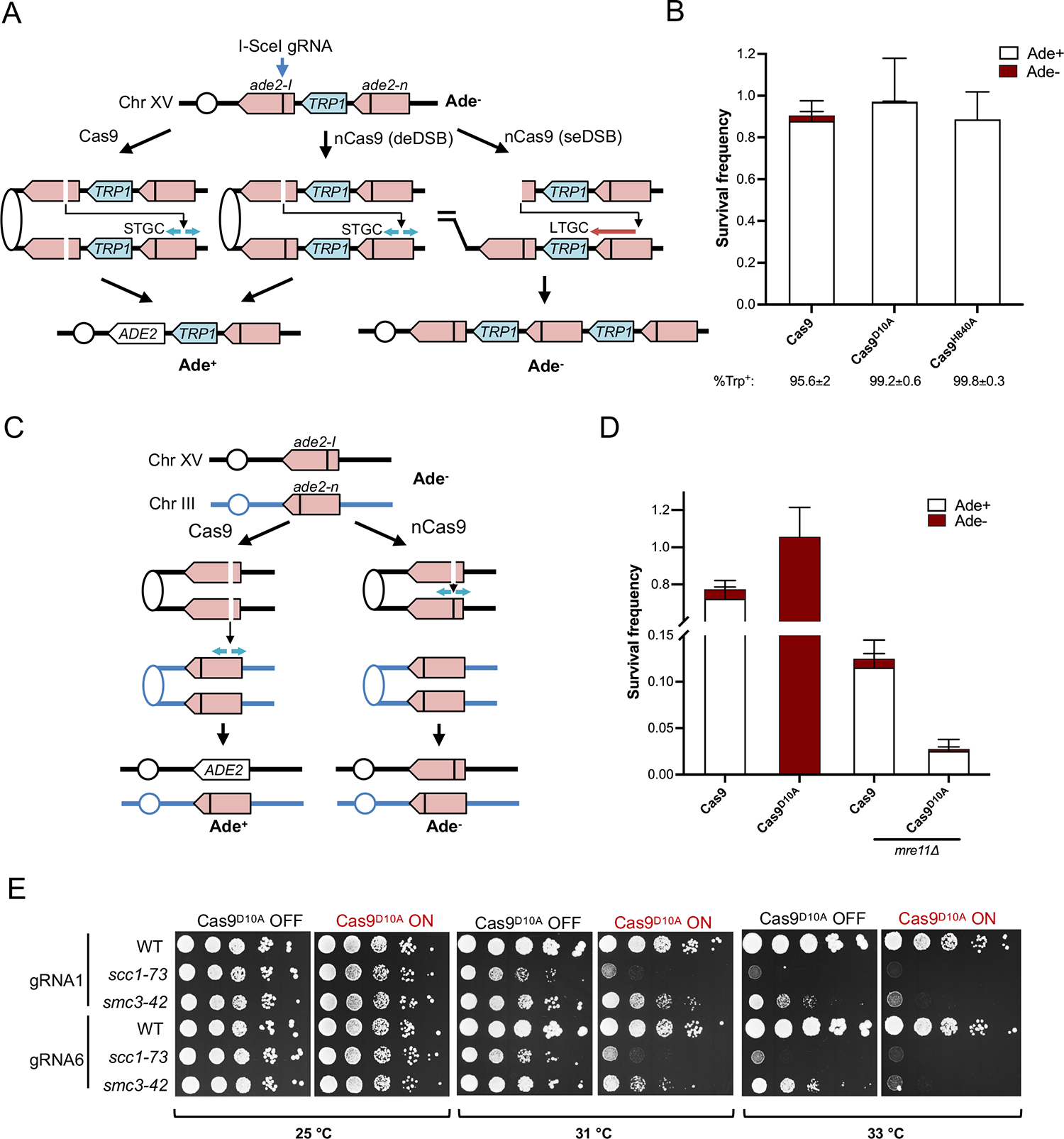
Cas9^D10A^-induced DSBs are preferentially repaired by sister-chromatid recombination. A. Schematic of the direct-repeat recombination reporter and expected outcomes of Cas9 or nCas9-induced recombination. A gRNA was designed to target the I-SceI cut site inserted at the AatII site within one copy of *ade2* and the other allele (*ade2-n*) has a frameshift mutation at the NdeI site ^42^. The sites of the mutations are indicated by vertical bars. Repair of a deDSB by short tract gene conversion (STGC) would generate an Ade^+^ recombinant, whereas long-tract gene conversion (LTGC) initiated from a seDSB would copy through the *ade2-n* mutation present in the donor giving rise to an Ade^-^ recombinant. B. Log-phase cells were plated on YPD or YPD+*β*-estradiol and percent survival following induction of Cas9/nCas9 was determined as described in Methods. The Ade phenotype was determined by white (Ade^+^) or red (Ade^-^) colony color and Trp phenotype by replica plating to medium lacking tryptophan. Error bars show the mean survival frequency from at least four trials of each strain. C. Schematic of the inter-chromosomal recombination assay and expected outcomes. D Log-phase cells were plated on YPD or YPD+*β*-estradiol and percent survival following induction of Cas9/Cas9^D10A^ was determined as described in Methods. The Ade phenotype was determined by white (Ade^+^) or red (Ade^-^) colony color. Error bars show the mean survival frequency from four trials of each strain. E. Sensitivity of WT, *scc1-73* and *smc3-42* strains to expression of Cas9^D10A^ paired with gRNA1 or gRNA6. Ten-fold serial dilutions of the strains were spotted on medium +/- *β*-estradiol and grown for 3 days at the indicated temperatures.

Since the survival assays indicate efficient repair of a replication-dependent DSB using the sister chromatid, we anticipated that a nick would not induce recombination between non-sister chromatids. To test this idea, we used an inter-chromosomal assay that consists of the same *ade2-I* allele on Chr XV, but the *ade2-n* repair template is integrated at the *LEU2* locus on Chr III (Figure 3C).^37^ In this scenario, the Cas9^D10A^-induced DSB has the potential to repair from the sister chromatid, resulting in an Ade^-^ event. However, for the fork collapse to result in an Ade^+^ event, the ectopic *ade2-n* donor must be used (Figure 3C). We found that expression of Cas9^D10A^ and I-SceI gRNA failed to stimulate Ade^+^ recombinants (Figure 3D). By contrast, induction of Cas9 resulted in reduced cell survival (77%), but most of the surviving colonies were Ade^+^, similar to the outcome from an I-SceI-induced DSB^37^. We presume that the strong preference for Ade^-^ repair outcomes with Cas9 nickase is due to cohesion of sister chromatids, a process in which the MRX complex has been implicated.^45–49^ Therefore, we reasoned that eliminating Mre11 may release the constraint on sister-chromatid recombination and result in Ade^+^ events from ectopic recombination. Indeed, when we deleted *MRE11*, we found that most of the survivors were Ade^+^ (Figure 3D). There was a notable drop in survival in *mre11Δ* cells, emphasizing the importance of Mre11 for efficient repair of a collapsed fork (Figure 3D). For the Cas9-induced DSB, the reduced survival of WT cells was further exacerbated by deletion of *MRE11* (Figure 3D). It is notable that in contrast to WT cells, *mre11Δ* cells show a lower survival to a nick than a DSB, despite the presence of an ectopic repair template (Figure 3D). These data indicate that a nick-induced fork collapse is preferentially repaired by sister chromatid recombination and that Mre11 plays an important role in reinforcing this preference.

### Role of sister chromatid cohesion in repair of collapsed replication forks

To further address the role of sister chromatid cohesion in repair of collapsed replication forks, we analyzed two temperature-sensitive mutants of the cohesin complex, *scc1-73* and *smc3-42*. At the semi-permissive temperature of 31 °C for *scc1-73* or 33 °C for *smc3-42*, expression of Cas9^D10A^ with gRNA1 or gRNA6 resulted in growth inhibition (Figure 3E), in line with a previous study showing that cohesin is partially required for HO nick-induced sister-chromatid recombination.^12^ We also tested the requirement for *CTF4* and *CTF8*, two non-essential genes that reinforce cohesion establishment during S-phase.^50,51^ The *ctf4Δ* mutant showed high sensitivity to Cas9^D10A^ expression, particularly with gRNA6, whereas *ctf8Δ* exhibited a modest but reproducible growth inhibition with both gRNAs (Figure S3B).

### A genome-wide screen identifies dependencies for replication-associated DSB repair

After identifying several factors necessary for repair of nickase-induced DNA damage, we next sought to identify other factors/pathways in an unbiased manner by performing a genome-wide screen for mutants that are sensitive to induction of Cas9^D10A^. We used selective ploidy ablation (SPA) and plasmid transfer to introduce plasmids containing Cas9^D10A^ and a gRNA into the yeast deletion strain collection (see Methods for details).^52^ Since, the deletion strain collection is a different genetic background to our standard laboratory strain (W303 background), we first verified that S288C derivatives deleted for *MRE11* or *RAD51* show sensitivity to nick induction (Figure S4A). For the screen, we used four plasmids: three expressed Cas9^D10A^ with gRNA1, gRNA6, or a scrambled gRNA1, and one was an empty vector, the latter two serving as controls (Figure S4B). After carrying out the SPA procedure, colony size was assessed, and a reduction in colony size was indicative of a genetic dependency. Two repeats of the screen were performed. Overall, the strains exhibiting growth defects were consistent between the two repeats for both gRNA1 and gRNA6 (Figure S4C).

The strains that exhibited a significant growth reduction were overwhelmingly mutants of the *RAD52* epistasis group (Figure 4A). One notable exception is the lack of *RAD59* from either gRNA group. This finding differs from a previous study using the HO-nick TINV system,^11^ but may be explained by the possibility of repair by BIR and single-strand annealing between short homologies using an intrachromosomal template in the plasmid-based system. We also failed to recover *SAE2* from the screen. The lack of sensitivity in the *sae2Δ* mutant is consistent with our finding that the *mre11-H125N* strain did not show growth sensitivity after expression of Cas9^D10A^ with either gRNA (Figure S3), confirming that Mre11 nuclease activity is not a requirement for collapsed fork repair.

**Figure 4.**
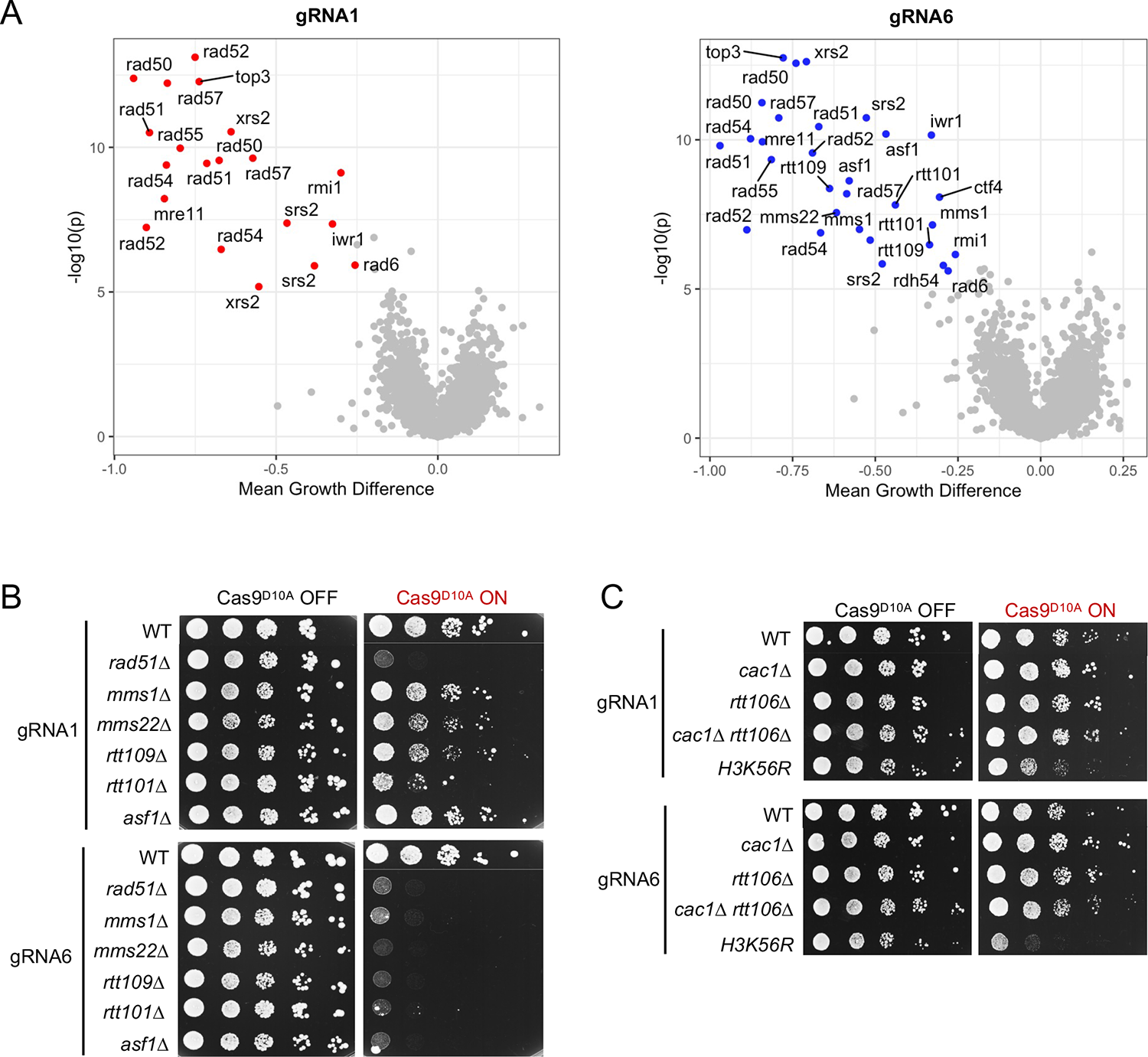
Genome-wide screen for mutants defective in the repair of broken replication forks. A. Mean difference in colony size between each gRNA and scrambled gRNA vs -log10(P-value) is represented for gRNA1 and gRNA6. Plots show combined data for two repeats of the screen using the non-essential deletion library. Colored points (gRNA1=red; gRNA6=blue) meet a cutoff of a mean growth difference of less than −0.25 or greater than 0.25 and -log10(p) > 5. Gene names of those showing a significant growth difference are included. B Spot assays to assess cell viability after expression of Cas9^D10A^ with gRNA1 or gRNA6 in the indicated strain backgrounds. C. Spot assays to assess cell viability after expression of Cas9^D10A^ with gRNA1 or gRNA6 in the indicated strain backgrounds. Ten-fold serial dilutions of log phase cultures were spotted on synthetic media lacking leucine either with (Cas9^D10A^ ON) or without (Cas9^D10A^ OFF) β-estradiol.

Several other hits for both gRNAs, including *SRS2, TOP3* and *RMI1*, may be involved in processing of the D-loop or HJs that arise during repair. Srs2 is an anti-recombinase, whose function may be important for disrupting non-sister strand invasion events or disrupting the invasion of ssDNA behind the migrating D-loop, as has been found in BIR.^53–56^ *TOP3* and *RMI1*, but not *SGS1*, were identified in the screen, consistent with our finding that the *sgs1Δ* mutant is not sensitive to Cas9^D10A^-induced nicking (Figure S3). The Top3-Rmi1 complex can disrupt D-loops, decatenate double HJ intermediates and resolve chromosome entanglements.^57–59^ which may arise during repair of the collapsed replication fork. *RAD6* was also a weak hit with both gRNAs. Rad6 is a ubiquitin conjugating enzyme that forms complexes with several other proteins, including Rad18 and Bre1. The Rad6-Rad18 complex monoubiquitylates proliferating cell nuclear antigen (PCNA) to promote translesion synthesis (TLS).^60–62^ The Rad6-Bre1 complex monoubiquitylates H2B,^63,64^ a function recently linked to sister chromatid cohesion.^65^ Since none of Rad6’s partners appeared as hits in the screen, additional testing will be needed to determine which function of Rad6 is important for its resistance to nCas9 expression.

We did not recover mutants defective for the DNA damage checkpoint in the screen. Consistent with this finding, we detected only minimal phosphorylation of Rad53 after induction of Cas9^D10A^ with gRNA1 or gRNA6 in asynchronous cultures (Figure S5). Elimination of Rad51 resulted in a strong checkpoint response, as expected when DNA repair is compromised. Additionally, a poorly repaired DSB generated by expression of Cas9 in WT cells resulted in robust Rad53 phosphorylation (Figure S5). Thus, the failure to detect Rad53 activation when nCas9 is induced is likely because the replication-dependent DSB is rapidly repaired without generation of extensive tracts of ssDNA and is not due to nCas9 shielding the ends from resection and checkpoint activation.

Interestingly, we recovered a group of mutants that were specifically sensitive to expression of Cas9^D10A^ with gRNA6, but not gRNA1 (Figure 4A). These include members of the RCNA pathway: *rtt109Δ, mms22Δ, asf1Δ, rtt101Δ*, and *mms1Δ.* Previous studies have shown that components of the RCNA pathway are required for resistance to CPT, MMS and unscheduled RNaseH2 expression, and for MMS-induced sister chromatid recombination.^21–23,66^ In this pathway, Asf1 acts as a histone H3-H4 chaperone, presenting the dimer to the acetyltransferase Rtt109, which acetylates H3 on lysine 56.^67–69^ H3K56ac increases affinity of the H3-H4 dimer for Rtt101-Mms1-Mms22, which ubiquitylates H3 at several locations (H3K121, K122, K125), causing the release of Asf1 and increasing affinity for the histone deposition machinery, CAF-1 and Rtt106.^70,71^ CAF-1 and Rtt106 deposit H3K56ac-H4 behind the replication fork.^72^ *CTF4* was also identified as a gRNA6-specific hit and it is noteworthy that Ctf4 interacts with the Rtt101-Mms1-Mms22 complex, which has been implicated in establishing H3K56ac and regulating the replisome in response to replication stress.^73–75^ However, the gRNA6 specificity of the *ctf4Δ* mutant is less robust than the RCNA mutants and it just failed to meet the cut off in the screen with gRNA1 (see Figure S3B).

To explore the discrepancy between gRNA1 and gRNA6 in the RCNA mutants, we first validated the sensitivities individually by transforming the Cas9^D10A^-gRNA1 and Cas9^D10A^-gRNA6 plasmids into each of these mutant strains in the W303 genetic background and testing for growth inhibition by spot assay (Figure 4B). These results confirmed the increased sensitivity with Cas9^D10A^-gRNA6 observed in the screen. As a comparison, the *rad51Δ* mutant is highly sensitive to expression of both gRNAs, indicating that the differential effect we observe for RCNA mutants is specific to this group of genes. Since the RCNA pathway is responsible for acetylating lysine 56 on H3, we tested an *H3K56R* acetylation deficient mutant and observed growth inhibition with gRNA6, confirming that it is the failure to introduce this histone modification that is responsible for the sensitivity observed (Figure 4C). CAF-1 and Rtt106 cooperate to deposit H3K56ac behind forks and the *cac1Δ rtt106Δ* double mutant was previously shown to be sensitive to genotoxic agents.^76^ However, we did not observe sensitivity to Cas9^D10A^ expression with gRNA6 in this mutant (Figure 4C). One possible explanation is that other H3-H4 chaperones compensate for loss of CAF-1 and Rtt106. Consistent with this idea, incorporation of H3K56Ac into chromatin is reduced but not abolished in the *cac1Δ rtt106Δ* double mutant.^76^ Furthermore, the *cac1Δ rtt106Δ* shows modest sensitivity to low doses of DNA damaging agents and is consistently less sensitive than other RCNA mutants, like *rtt109Δ.* ^76,77^

### The RCNA pathway is specifically required for repair of a leading strand fork collapse

We reasoned that the differential requirement for the RCNA pathway could be due to the location of the nicks induced by gRNA1 and gRNA6. Cas9^D10A^ with gRNA1 nicks the lagging strand template whereas Cas9^D10A^-gRNA6 nicks the leading strand template. To determine whether the RCNA pathway is specifically required to repair a leading strand fork collapse, we tested sensitivity of *mms22Δ* and *rtt109Δ* mutants to Cas9^D10A^ expression using gRNA14 or gRNA20, which cleave the opposite strand to gRNA1 or gRNA6, respectively (Figure 5A and Figure S1). Interestingly, the *mms22Δ* and *rtt109Δ* mutants displayed high sensitivity to nicking by gRNA14, but less sensitivity to gRNA20, the reverse of the growth defects observed for gRNA1 and gRNA6 (Figure 5A). By contrast, growth of the *rad51Δ* mutant was inhibited by Cas9^D10A^ with all gRNAs tested. The same trend of higher sensitivity to a leading strand template nick was observed with Cas9^H840A^ but to a lesser extent than found for Cas9^D10A^ (Figure S6).

**Figure 5.**
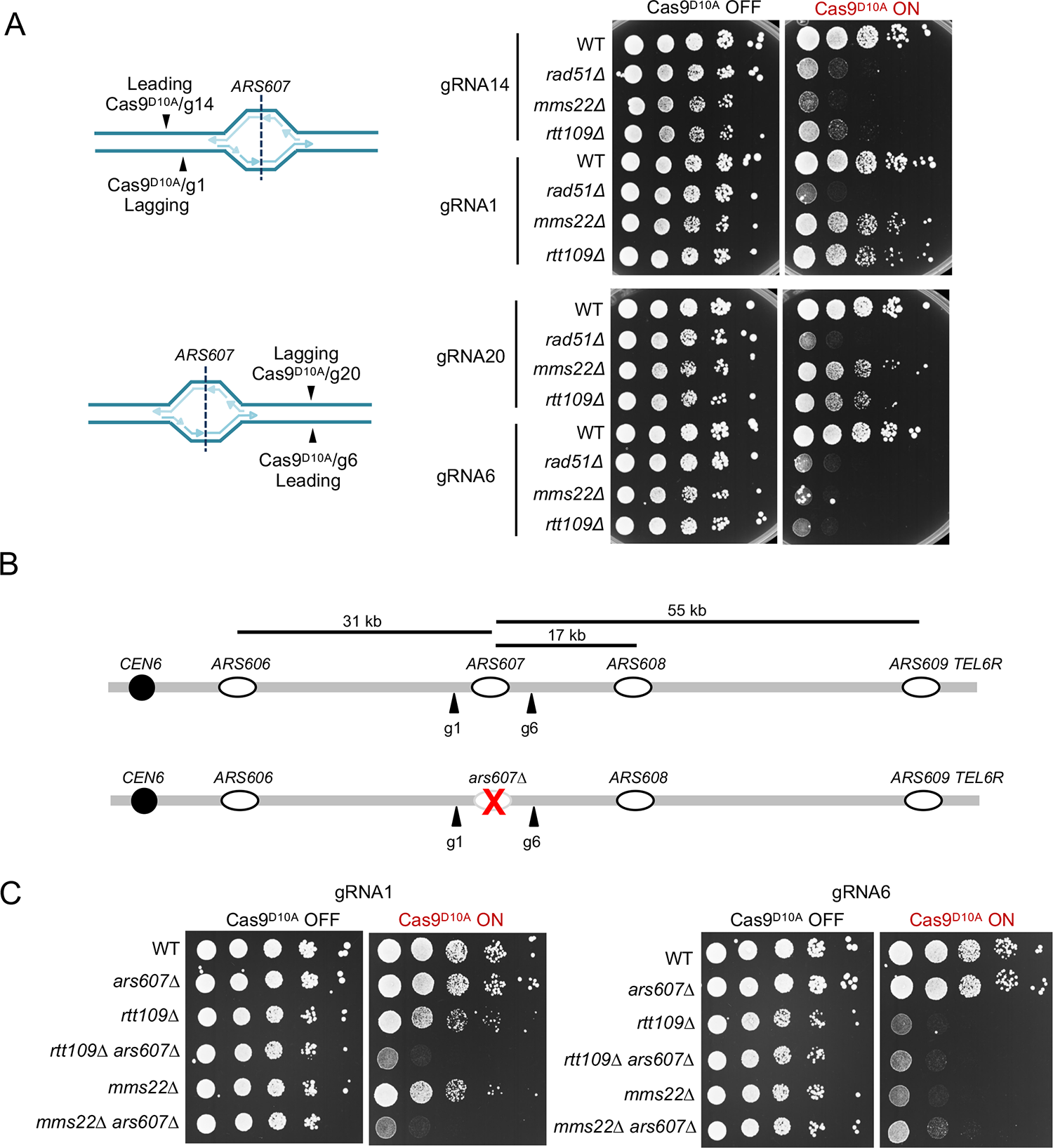
The RCNA pathway is required for repair of a leading strand fork collapse. A. Schematic showing which strand is cleaved by the indicated gRNAs with Cas9^D10A^ (left) and corresponding spot assays (right). Ten-fold serial dilutions of the indicated strains were spotted on synthetic media lacking leucine either with (Cas9^D10A^ ON) or without (Cas9^D10A^ OFF) β-estradiol. B. Schematic showing the locations of replication origins on the right arm of Chr. VI relative to gRNA1 and gRNA6. Deletion of *ARS607* is expected to reverse the strands cleaved by Cas9^D10A^ with gRNA1 since *ARS606* is more efficient and fires earlier than *ARS608* and *ARS609*. C. Ten-fold serial dilutions of the indicated strains were spotted on synthetic media lacking leucine either with (Cas9^D10A^ ON) or without (Cas9^D10A^ OFF) β-estradiol.

To further test the template specificity for the RCNA pathway, we deleted *ARS607* so that the nick generated by Cas9^D10A^-gRNA1 would be on the leading strand template from a replication fork initiated at *ARS606*, the same template strand as the nick generated by Cas9^D10A^-gRNA6 (Figure 5B). *ARS608* is a very inefficient origin and *ARS609* is late firing ^26,27^; thus, leftward moving forks originating from these origins would not be expected to contribute to replication of the gRNA1 target sequence. Deletion of *ARS607* alone did not affect sensitivity of the WT strain with either guide RNA (Figure 5C). However, expression of Cas9^D10A^-gRNA1 was lethal in *ars607Δ mms22Δ* and *ars607Δ rtt109Δ* strains (Figure 5C), consistent with an increased need for the RCNA pathway to repair a leading strand fork collapse. We observed a slight growth restoration of the *ars607Δ mms22Δ* and *ars607Δ rtt109Δ*strains to Cas9^D10A^-gRNA6 expression, suggesting that some fraction of replication forks reaching the gRNA6-induced nick initiate from *ARS608* or *ARS609* in the absence of *ARS607*, creating a deDSB on the lagging strand.

## DISCUSSION

Here we have established a system in to induce nicks on either the leading or lagging strand template that are converted to DSBs during S-phase. We show that Rad51-dependent homologous recombination is essential for repair of these lesions and there is a strong bias to repair using a sister chromatid template. Although the core HR factors, HJ resolvases and cohesin components are essential to repair nicks induced on the leading and lagging strand templates, we identify a unique requirement for the RCNA pathway to repair lesions induced on the leading strand template. These findings are discussed in more detail below.

By physical methods, we show that Cas9^D10A^ induces S-phase dependent DSBs. Contrary to expectation,^5^ we observe deDSBs for nicks induced on the lagging strand template. Since deletion of neighboring origins does not prevent deDSB formation, we favor the hypothesis that CMG continues unwinding on the leading strand template at a Cas9^D10A^-induced nick,^78^ displacing the lagging strand for initiation of an Okazaki fragment that would extend to the nick site. This finding agrees with recent observations in human cells using the Cas9^D10A^ nickase and studies using a reconstituted *Xenopus laevis* egg extract system showing that the MCM complex and CDC45 are retained on chromatin following replication fork collapse.^79,80^ The Cas9^D10A^ post-cleavage complex might be permissive for CMG to continue translocation on the leading strand instead of translocating onto duplex DNA, as has been reported for a reconstituted system.^5^ The recombination products generated by Flp-mediated nicking in mouse cells are also consistent with repair of a replication-dependent deDSB.^10^ Surprisingly, we observe a deDSB with one of the gRNAs that targets the leading strand. We note that deDSBs are also detected using the HO-nick and Flp-nick systems in a plasmid context in budding yeast, and in the *S. pombe* genome using gpII and nCas9 nickases.^11,81–83^ Although we still observed a deDSB when *ARS608* and *ARS609* were deleted, we cannot rule out fork convergence from other poorly annotated origins in this reqion of the chromosome that are activated by fork collapse.^32^ Generation of a replication-dependent deDSB instead of seDSB has advantages for maintaining genome integrity. If a fork collapse were to generate a seDSB, ligation of the hybrid nick formed between the parental duplex and nascent strand could allow local re-replication from an adjacent origin.^84^ Such an event could be prevented if the hybrid nick were left unligated, but strand invasion into the sister chromatid template to restart replication might then be compromised. Bypass of the nick and continued synthesis to generate a deDSB would avoid this scenario. In addition, extensive DNA synthesis from a seDSB is more mutagenic than normal S-phase synthesis, contributing to genomic instability.^85^

Regardless of whether a seDSB or a deDSB is formed, we find that repair is highly dependent on HR factors and is independent of NHEJ (Figure 6). A replication-dependent deDSB formed by nCas9 may be inefficiently repaired by end joining because one end would have an overhang up to the size of an Okazaki fragment rendering it a poor substrate for canonical NHEJ. Yeast cells lack Pol *θ*-mediated end joining (TMEJ) and repair using microhomologies is inefficient.^86^ In agreement with our findings, nCas9 and Flp-nick-induced HR in murine and human cells is not suppressed by canonical NHEJ factors in contrast to Cas9-induced events.^10,17^ Rare end-joined products can be detected in response to nCas9 expression in mammalian cells and these exhibit deletions with microhomologies at the junctions, a hallmark of TMEJ. Interestingly, end-joined products are increased in the absence of BRCA1, consistent with use of alternate error-prone repair pathways in the absence of HR.^16^ Surprisingly, we observe a requirement for HJ resolvases, but not for Sgs1-mediated dHJ dissolution, in tolerance of nicks induced by Cas9^D10A^. This finding could indicate that single HJs are generated during repair, either by D-loop processing or fork convergence.

**Figure 6.**
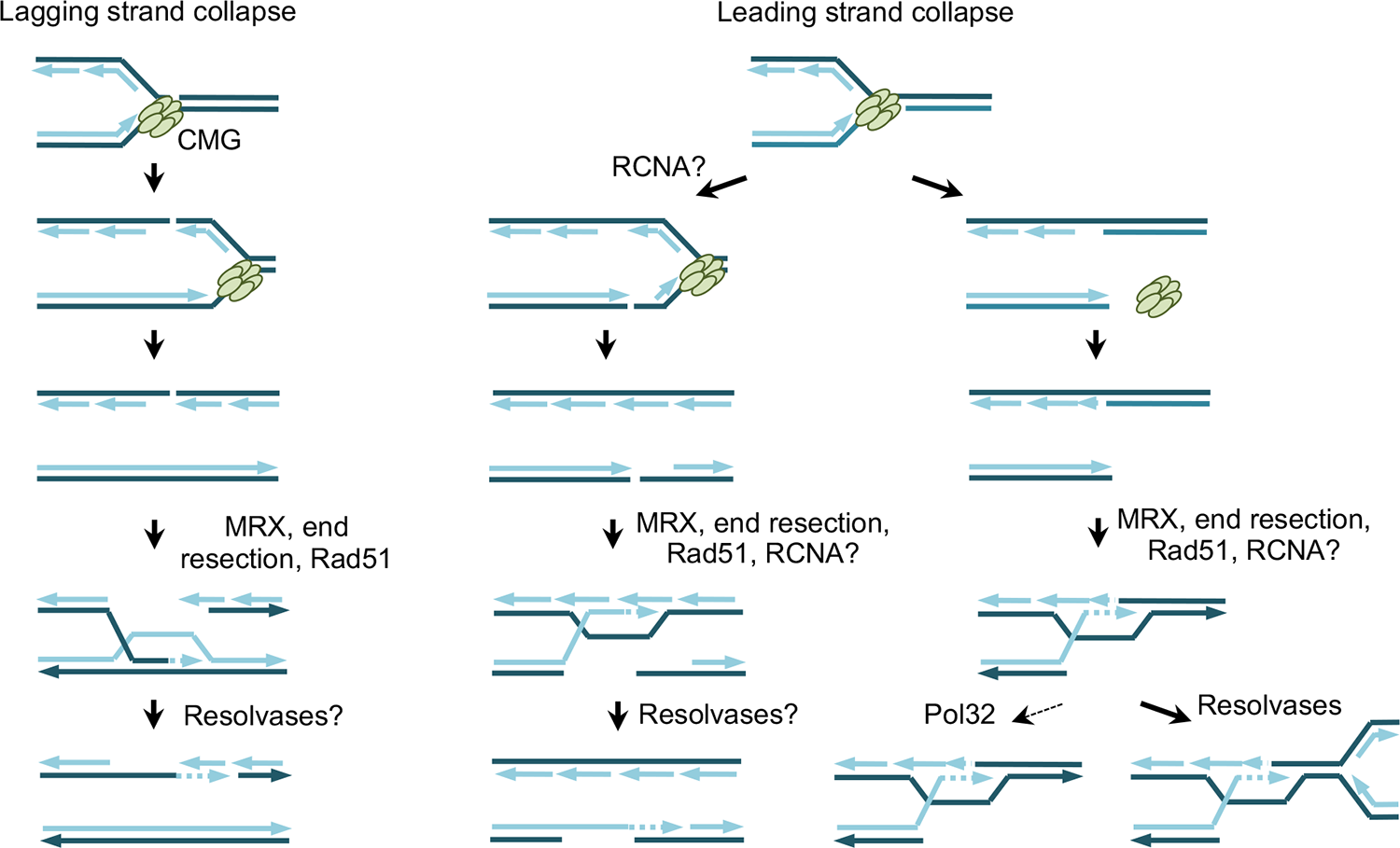
Model for the repair of replication-dependent DSBs. Replication fork collapse at nicks induced by Cas9^D10A^ on the leading and lagging strand templates can generate single-end or deDSBs, both of which require the MRX complex, long-range end resection and Rad51-dependent strand invasion of the sister chromatid for repair. The RCNA pathway is more important for repair of replication-dependent DSBs on the leading strand template. This requirement could be due to RCNA components stabilizing the replisome after encounter with a leading strand nick, facilitating invasion of the lagging strand template, or a post-invasion role in establishing chromatin on the leading strand. The requirement for HJ resolvases suggests that second end capture at the deDSB involves processing of branched DNA structures that cannot be removed by the Sgs1-Top3-Rmi1 dHJ dissolution complex. The minor role of Pol32 indicates that extensive DNA synthesis within the context of a D-loop is dispensable for repair of replication-dependent DSBs.

We found that nCas9-induced nicks at the *ade2-I* allele of an *ade2* direct repeat reporter stimulate recombination with the non-allelic donor located 4.3 kb downstream of *ade2-I*. Most products result from STGC, regardless of whether the nick is on the leading or lagging strand template, consistent with repair of a deDSB. Although we did not identify products consistent with LTGC/BIR, studies in mammalian cells have shown an increased LTGC/STGC ratio in response to nCas9 or Flp-nick expression.^10,16,19^ These findings could be explained by BIR initiating at a seDSB or to more extensive DNA synthesis associated with HR during S-phase in mammalian cells.

By contrast, very few recombinants were generated by Cas9^D10A^ expression when the donor was located on a different chromosome, consistent with a strong preference to repair using a sister chromatid template. Cas9 does stimulate inter-chromosomal recombination indicating that repair can default to an ectopic donor in the absence of an intact sister chromatid. While deletion of *MRE11* allowed for inter-chromosomal recombination, it also reduced the survival frequency. We assume that deletion of *MRE11* allows for inter-chromosomal recombination due to a loss of sister chromatid tethering by the MRX complex, as has been suggested previously,^45–48,87^ or indirectly via diminished recruitment of cohesin.^49,88,89^ Notably, the *scc1-73* and *smc3-42* conditional alleles confer high sensitivity to nCas9 expression, consistent with the need to tether sister chromatids for efficient repair. The observation that in WT cells, nearly all nick-induced repair events were Ade^-^, while nearly all Cas9-induced repair events were Ade^+^ highlights the propensity for replication-associated DSBs to repair by sister-chromatid recombination. Our findings are consistent with studies in human cells showing a single nick induced by nCas9 fails to induce inter-homolog recombination.^90^ However, Cas9 nickases can induce allelic and ectopic inter-chromosomal gene conversion in Drosophila somatic cells,^91^ and Flp-nick is reported to induce allelic recombination in yeast.^92^ More studies will be needed to determine how well nicks can induce allelic recombination in different systems and genetic control of these events.

The genome-wide screen was carried out with the intent of identifying novel factors required for the repair of replication-dependent DSBs. Since CPT can induce replication fork collapse, factors identified in screens for CPT sensitivity should largely overlap with factors that are sensitive to nCas9 expression. First and foremost, the *RAD52* epistasis group is overwhelmingly represented in the nCas9 screen, which is consistent with results from CPT sensitivity screens, Flp-nick sensitivity, and HO-nick induced sister-chromatid exchange.^4,8,11,21,22^ There are some genes, for example, *MUS81* and *MMS4*, that show a requirement for survival to CPT but not with nCas9.^21,22^ It is only when *YEN1* is eliminated in the *mus81Δ* background that we observe growth inhibition (Figure 2D). This finding may be related to the number of structures induced with CPT that would require the action of Mus81-Mms4, compared to the single lesion induced by nCas9. Indeed, at lower doses of CPT, the growth defect of *mus81Δ* cells is minimal, but *mus81Δ yen1Δ* cells are inviable.^93^ Likewise, sensitivity to the single lesion induced by Flp-nick required deletion of both *MUS81* and *YEN1* ^4^. Since CPT also induces fork reversal,^94^ another possibility is that some mutants identified in CPT sensitivity screens could be defective for generation or processing of reversed forks instead of, or in addition to, repair of broken forks.

One of the more interesting findings from the screen was the gRNA specific sensitivity of RCNA mutants, a phenotype not observed for other mutants tested, such as *rad51Δ*. The RCNA pathway has long been implicated in resistance to replication stress and DNA damage-induced sister-chromatid recombination;^21–23,66,95–99^ however, the exact role of these proteins in the replication stress response is not completely understood. Our data indicate a specific requirement for the RCNA factors to repair a leading strand fork collapse generated by Cas9^D10A^, whereas a lagging strand collapse is tolerated. This bias is still observed but to a lesser extent using Cas9^H840A^. These requirements do not necessarily fit with the Rad51 loading role of Mms22^100, 101^ because Rad51 loading should be important regardless of whether the replication-associated DSB is on the leading or lagging strand template. Similarly, the sister-chromatid cohesion function of RCNA^102^ is unlikely to contribute to the gRNA specificity since conditional mutants of the cohesin complex showed equal sensitivity to gRNA1 and gRNA6.

The strand bias of the RCNA mutants with Cas9^D10A^ could be due to either the nature of the fork collapse or the mechanism of repair. Components of the RCNA pathway have been shown to stabilize replisomes during replication stress and could be important to maintain the replisome after encountering the nick, or for efficient convergence from neighboring replication forks.^77,103,104^ A collapse on the leading strand would require invasion of the lagging strand template to initiate replication restart (Figure 6). The lagging strand template could be less favorable for strand invasion if Okazaki fragments are not completely ligated or if there is an imbalance of nucleosomes. Okazaki fragment lengths are not noticeably different in RCNA mutants but changes in ligation efficiency have not been evaluated.^105^ There is a slight bias for incorporation of H3K56Ac-containing nucleosomes on the leading strand template, whereas parental nucleosomes are preferentially transferred to the lagging strand template.^106^ This bias, which might be accentuated in the RCNA mutants, could make the lagging strand less accessible for strand invasion. The H3K56Ac mark has been suggested to loosen the histone-DNA interaction at the entry and exit points of the nucleosome, which may allow for a more favorable chromatin environment for strand invasion on the leading strand.^68^

Finally, the role of RCNA in sister chromatid recombination could be post strand invasion. Nucleosomes displaced by end resection would need to be replaced after repair synthesis and this would require incorporation of newly synthesized histones. A post-strand invasion role for RCNA in restoration of chromatin structure at DSBs has been proposed previously and linked to DNA damage checkpoint adaptation.^107^ Although restoration of nucleosomes post repair should be important for leading and lagging strand collapses, if the nucleosome composition of the leading and lagging strand templates is different, then chromatin restoration on the leading strand might be more challenging in the absence of the RCNA pathway.

### LIMITATIONS OF THE STUDY

The use of nCas9 to model replication fork collapse has many attractive features, including site and strand specificity, but may not fully mimic spontaneous lesions. Cas9 is known to remain associated with the target sequence after cleavage and this could impact how the replisome behaves when encountering a nick. Indeed, the deDSB observed using Cas9^D10A^ to generate a nick on the lagging strand template could be specific to use of this nickase since Cas9^H840A^ has been shown to generate seDSBs at both leading and lagging strand nicks in a cell-free system. ^5^ Furthermore, the RNA-DNA hybrid formed during Cas9 cleavage could affect nick recognition and processing. Another limitation is that little is known about the dynamic interaction between nCas9 and the target sequence. In our studies, the nickase is continually expressed in cells after induction and may undergo multiple cycles of binding, nicking and release. We imagine that many of the nicks are ligated prior to arrival of a replication fork. For nicks that result in fork collapse, nCas9 could also nick the sister chromatid that is used as a template for strand invasion, potentially influencing repair dynamics. Despite these limitations, the genetic requirements we observe for repair of nCas9-induced DNA damage closely match those reported with other nickases and CPT-induced lesion,^4,8,9,11,21,22,99^ suggesting that the repair mechanisms used are common to SSBs and are not specific to nCas9.

## METHODS

### Media and yeast strains

Complete yeast media contained 1% yeast extract, 2% peptone, 10 µg/mL adenine, and 2% glucose (YPAD) as a carbon source. Synthetic media contained 1X yeast nitrogen base, 1X amino acid dropout mix, and 2% glucose. For *β*-estradiol induction of (n)Cas9, a 10 mM stock of *β*-estradiol was diluted and spread on plates or added to liquid media for a final concentration of 2 µM.

All yeast strains are in the *RAD5*-corrected W303 background (*leu2-3*,*112 trp1-1 can1-100 ura3-1 ade2-1 his3-11,15*) unless otherwise noted and are listed in Supplementary Table 1.

The BY4742/S288C strain background (*MATα his3Δ1 leu2Δ0 lys2Δ0 ura3Δ0*) was used for the library screen and for the spot assays shown in Figure S4A. Strains were constructed by standard genetic methods. Lithium acetate transformation was used to introduce deletion cassettes containing a marker of choice and homology arms flanking the gene to be deleted, or episomal plasmids. Other strains were made by genetic cross, followed by tetrad dissection and marker selection.

### Cas9 and gRNA plasmids

#### Integrating Cas9 plasmids

Integrating plasmids were constructed in pRG20xMX backbones and integrated into the genome as previously described.^108^ A plasmid containing LexO-Cas9-NLS-3xFLAG-TCYC1 PACT1-LexA-ER_LBD-B112-TCYC1 in a pRG203MX (*HIS3*) plasmid (pLS504) was used for all experiments with integrated Cas9 (Supplementary Table 3).^25^ Cas9^D10A^ was generated by site-directed mutagenesis of pLS504 using the QuikChange II XL Site-Directed Mutagenesis Kit (Agilent #200521) using primers MK11 and MK12, yielding plasmid pLS503. Cas9^H840A^ was made by site-directed mutagenesis of pLS504 using the Q5 site-directed mutagenesis kit (New England Biolabs #E0554S) using primers MK118 and MK119 to yield pLS517. The same procedure was used to introduce the H840A mutation into pLS503 (Cas9^D10A^) to yield the catalytically dead Cas9 (dCas9) plasmid (pLS640).

In a second set of plasmids, the Cas9 terminator (TCYC1) was switched to an ADH1 terminator (TADH1) by Gibson assembly using the HiFi DNA Assembly kit (New England Biolabs). Cas9^D10A^, Cas9^H840A^, and dCas9 were amplified from pLS503, pLS517, and pLS640, respectively, using primers MK249 and MK250. A previously constructed plasmid (pAA12)^25^ containing LexO-linker-TADH1 PACT1-LexA-ER_LBD-B112-TCYC1 was digested with ApaI and SacI within the linker region. The digested plasmid and amplified Cas9 fragments were assembled in a HiFi assembly reaction according to manufacturer’s protocol. This yielded pLS626 (Cas9^D10A^), pLS643 (dCas9), and pLS644 (Cas9^H840A^).

#### Integrating gRNA plasmids

gRNAs were expressed in a pRG205MX (LEU2) integrating plasmid backbone (pLS505). This plasmid contained a tyrosine tRNA promoter, HDV ribozyme, a sgRNA scaffold, and SNR52 RNA PolIII terminator (collectively referred to as the gRNA expression cassette), which was modified from a pCAS plasmid (Addgene # 60847) by replacing the gRNA cloning sequence with an XbaI and ZraI fragment by Robert Gnügge of this laboratory. The gRNA expression cassette was then cloned into the pRG205MX (pLS505) by Amr Al-Zain of this laboratory. An analogous plasmid was made in the pRG206MX (URA3) backbone (pLS545). For gRNA cloning, complementary oligos containing the gRNA sequence were designed to create a blunt end (ZraI compatible) on one side and a 5’ CTAG (XbaI compatible) overhang on the other. pLS505 was digested with XbaI and ZraI and gel purified. The gRNA-containing oligos were annealed and then ligated to the digested pLS505.

#### CEN plasmids for expression of Cas9 and gRNA

Centromeric plasmids were constructed that express both Cas9 and gRNA. For this purpose, the CEN plasmid pRS415 (LEU2) was used as the backbone, and Cas9-TADH1 and gRNAs were amplified from their respective integrating plasmids described above and assembled in a three-fragment HiFi reaction. pRS415 was linearized with HindIII, Cas9 constructs with the ADH1 terminator along with LexA transcription factor were amplified together using MK218 and MK221 primers, and gRNAs were amplified with MK219 and MK220 primers. PCR amplifications were performed using Phusion High-Fidelity PCR kit (New England Biolabs #M0530L) according to manufacturer’s protocol using 50 ng template DNA per 100µL reaction and 18 cycles. HiFi assembly reactions were carried out according to manufacturer’s protocol and assembly was confirmed by Sanger sequencing. A second set of plasmids using pRS414 (TRP1) was constructed using the same strategy, except pRS414 was linearized with ClaI. Cas9^D10A^-TADH1 was amplified from pLS626, Cas9^H840A^-TADH1 was amplified from pLS644, and dCas9 was amplified from pLS643. gRNA1 was amplified from pLS499, gRNA6 was amplified from pLS588, and scrambled gRNA was amplified from pLS631.

#### Physical analysis of DSBs

Unless otherwise noted in the figure legend, strains with integrated (n)Cas9 and gRNA cassettes were used for physical analysis of replication fork collapse. For asynchronous of Cas9 or Cas9^D10A^, single colonies of relevant strains were used to inoculate 5 mL YPAD cultures that were grown overnight at 30°C. The following day, cultures were diluted to an OD of 0.4 in YPAD and grown for 1 hour. 1 x 10^8^ cells were collected before induction (t0) and then *β*-estradiol was added to the remaining culture at a final concentration of 2 µM to induce Cas9/ Cas9^D10A^. The same number of cells were collected at each timepoint, sodium azide was added to 0.1% final, cells were collected by centrifugation, cell pellets were then washed with dH2O and stored on ice prior to embedding in agarose plugs. For cell synchronization, 50 ng/mL *α*-factor was added to *bar1Δ* derivatives at OD600 of 0.4 for 2 h to induce G1 arrest, nCas9 was then induced for 1 h (t-60).

Cells were collected, washed in fresh medium containing pronase and then released into S-phase. Cells were collected at 30 min time points and agarose plugs were prepared and processed as previously described.^109^ After digestion within plugs, the DNA was separated on a 0.8% agarose gel and then transferred to nylon membranes (Hybond N+). Membranes were hybridized with radiolabeled probes spanning the g1/g14 and g6/g20 recognition sites. Oligonucleotides used to generate probes by PCR amplification from genomic DNA are described in Supplementary Table 2.

### Spot assays for assessing growth inhibition by Cas9 or nCas9

For most of the spot assays, (n)Cas9 and the relevant gRNA were co-expressed from a centromeric plasmid. Single colonies were used to inoculate 2 mL SC-Leu or SC-Trp medium and cultures were grown overnight at 30°C while shaking. The following day, 10-fold serial dilutions were spotted on media with or without 2 µM *β*-estradiol. For strains that contained integrated Cas9 and gRNA, YPAD media was used for both liquid cultures and plates. Plates were grown for three days at 30°C, unless otherwise noted, after which plates were imaged with a desktop scanner.

### Recombination assays with (n)Cas9

Recombination assays using (n)Cas9 were performed similarly to those in which GAL-I-SceI was used to create the DSB in *ade2-I,*^37^ except for the following modifications. The strains used contained an integrated Cas9 or Cas9^D10A^ and integrated I-SceI gRNA. Single colonies were used to inoculate 2 mL YPAD cultures, which were incubated at 30°C while shaking for 4 hours. Cultures were then collected and diluted as described previously and plated on YPAD plates with or without 2 µM *β*-estradiol. Survival frequency and Ade^+^/Ade^-^ outcomes were determined as described previously.^37^ For the direct repeat assay, colonies were replica plated to SC-trp to score retention of the *TRP1* marker.

### Rad53 Western Blotting

Expression of Cas9/Cas9^D10A^ was induced in asynchronous cultures by addition of 2 µM *β*-estradiol. ∼1.4×10^8^ cells were removed at each timepoint and protein was extracted using trichloroacetic acid (TCA) precipitation and processed as described previously.^37^

### Genome-wide screen using selective ploidy ablation and Cas9-gRNA plasmid transfer

Selective ploidy ablation (SPA) and plasmid transfer takes advantage of a Universal Donor Strain (UDS), in which all 16 chromosomes contain a galactose promoter oriented toward the centromere and also contain a counter-selectable *URA3* marker ^52^. Briefly, a plasmid of interest can be transformed into the UDS, which can then be mated to a large number of target strains. After mating, galactose-induced transcription through the donor strain centromeres leads to their destabilization, leaving only the target strain chromosomes. Subsequent counter-selection of the remaining donor strain chromosomes with 5-fluoroorotic acid (5-FOA) ensures that surviving colonies only have the target strain chromosomes. This method has proven useful to introduce plasmids into yeast gene deletion libraries containing thousands of target strains when combined with automated pinning. For this screen, pRS415 (empty vector), pLS627 (Cas9^D10A^-gRNA1), pLS633 (Cas9^D10A^-gRNA6), and pLS632 (Cas9^D10A^-scr.gRNA) were transformed into the MAT*α* UDS (W8164-2C/ LSY5158-2C) for mating to the yeast gene deletion library. Donors and library strains were allowed to mate for 6 hours on YPD medium and then transferred to synthetic medium lacking leucine and containing galactose. These plates also contained 2 µM *β*-estradiol to induce expression of Cas9^D10A^. After 2 days of incubation colonies were transferred to synthetic medium lacking leucine and containing galactose, 5-FOA and 2 µM *β*-estradiol. Plates were then incubated for 3 days. All incubations were carried out at 30 °C. After the final incubation, plates were imaged at 300 dpi using a Scanmaker 9800XL-plus flatbed scanner (MicroTek) fitted with a TMA1600 transparency adapter.

### Analysis of genome-wide screen

Plate images were first processed using the *Screenmill* package for the R statistical programming language (https://github.com/robertjdreid/screenmill). This package annotates, calibrates and measures colony growth data. Raw colony growth data for each plate was normalized to median plate growth as described previously ^110^. For our purposes either pRS415 (EV) or pLS632 (Cas9^D10A^-scr.gRNA) can be used as the control strain from which to compare growth of strains with Cas9^D10A^ + gRNA1/6. Comparisons with the scrambled gRNA control are described below and used in the Results.

Using the normalized colony growth data, an average colony size of four replicates per strain, per plasmid was calculated. Average growth difference between Cas9^D10A^ + gRNA1/6 and Cas9^D10A^ + scr.gRNA was calculated for each strain, for two repeats of the screen. Data from both screens was combined and p-values were calculated using Student’s t test from the R *stats* package. Volcano plots represent the average growth difference (x-axis) between the scrambled gRNA conditions and the indicated gRNA vs. -log10(p-value) (y-axis). Correlation between the two repeats of the screen is visualized by plotting the growth difference of repeat 1 vs. repeat 2 for each gRNA.

## ACKNLOWEDGEMENTS

We thank J. Tercero, Z. Zhang, X. Zhao for gifts of yeast strains, and W.K. Holloman, A. Nicolas, and members of the Symington lab for review of the manuscript. We also thank A. Nussenzweig and R. Scully for communicating unpublished data. This work was supported by grants from the National Institutes of Health (R35 GM126997 to L.S.S.; R35 GM118180 to R.R. and T32 CA265828 to M.J.J.).

## AUTHOR CONTRIBUTIONS

MTK contributed to the experiments shown in Figures 1, 2, 3, 4, 5, S1, S2, S3, S4, S5 and S6; AS contributed to experiments shown in Figures 1, 5, S1, S2, and S6; MJJ contributed to experiments shown in Figures 2, 3, and S3. RJDR contributed to experiments shown in Figures 4 and S4. All authors contributed to study design, and MTK and LS wrote the manuscript.

---

**Figure S1.**
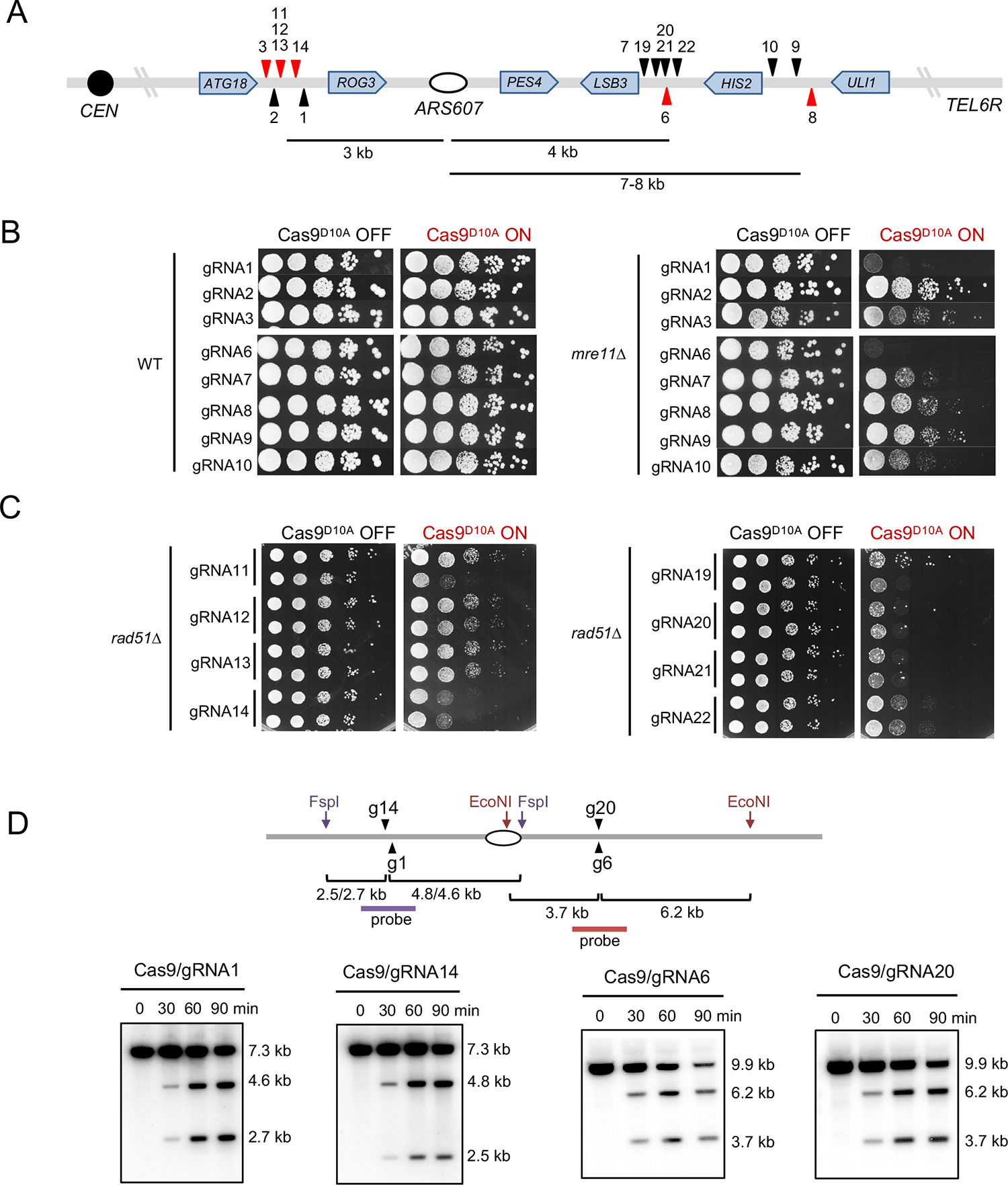
Efficient gRNAs were identified based on loss of viability in HR-deficient cells. A Schematic showing the location of gRNAs designed to target sequences centromere proximal and distal to *ARS607*. gRNAs targeting the leading strand are shown as red triangles and the gRNAs targeting the lagging strand are in black. All gRNAs were designed to target intergenic regions. B Ten-fold serial dilutions of WT or *mre11*Δ cells expressing Cas9^D10A^ and the indicated gRNAs were spotted on medium with or without β-estradiol. C Ten-fold serial dilutions of *rad51*Δ cells expressing Cas9^D10A^ and the indicated gRNAs were spotted on medium with or without β-estradiol (two independent transformants are shown for each gRNA). D. Schematic showing the relative positions of the gRNAs and restriction endonuclease cleavage sites. Southern blot analysis of WT strains with the indicated gRNAs prior to Cas9 induction and at 30 min intervals after addition of β−estradiol to the growth medium. Genomic DNA was digested FspI (g1 and g14) or EcoNI (g6 and g20) and the blots were hybridized with radioactive probes spanning the gRNA sites (purple or red lines, see Figure 1).

**Figure S2.**
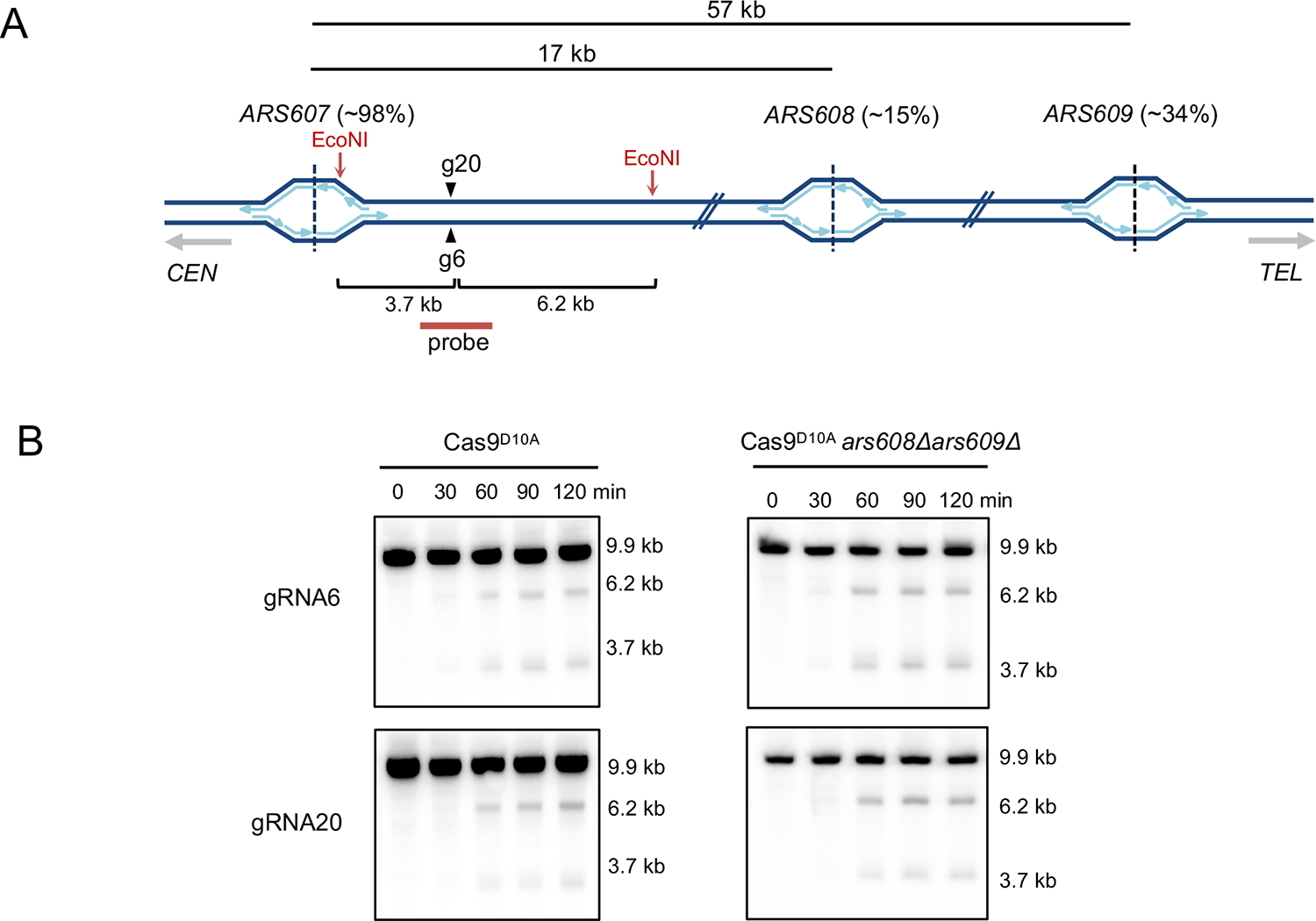
Double-ended DSBs are not the result of fork convergence. A. Schematic showing the location and efficiency or replication origins telomere distal to *ARS607*, the gRNA6 and gRNA20 sites (inverted triangles) and EcoNI site used for Southern blot analysis (see also Figure 1). B. Southern blot analysis of WT strains with the indicated gRNAs prior to Cas9^D10A^ induction and at 30 min intervals after addition of β-estradiol to the growth medium. Genomic DNA was digested with EcoNI and the blots were hybridized with radioactive probes spanning the gRNA sites.

**Figure S3.**
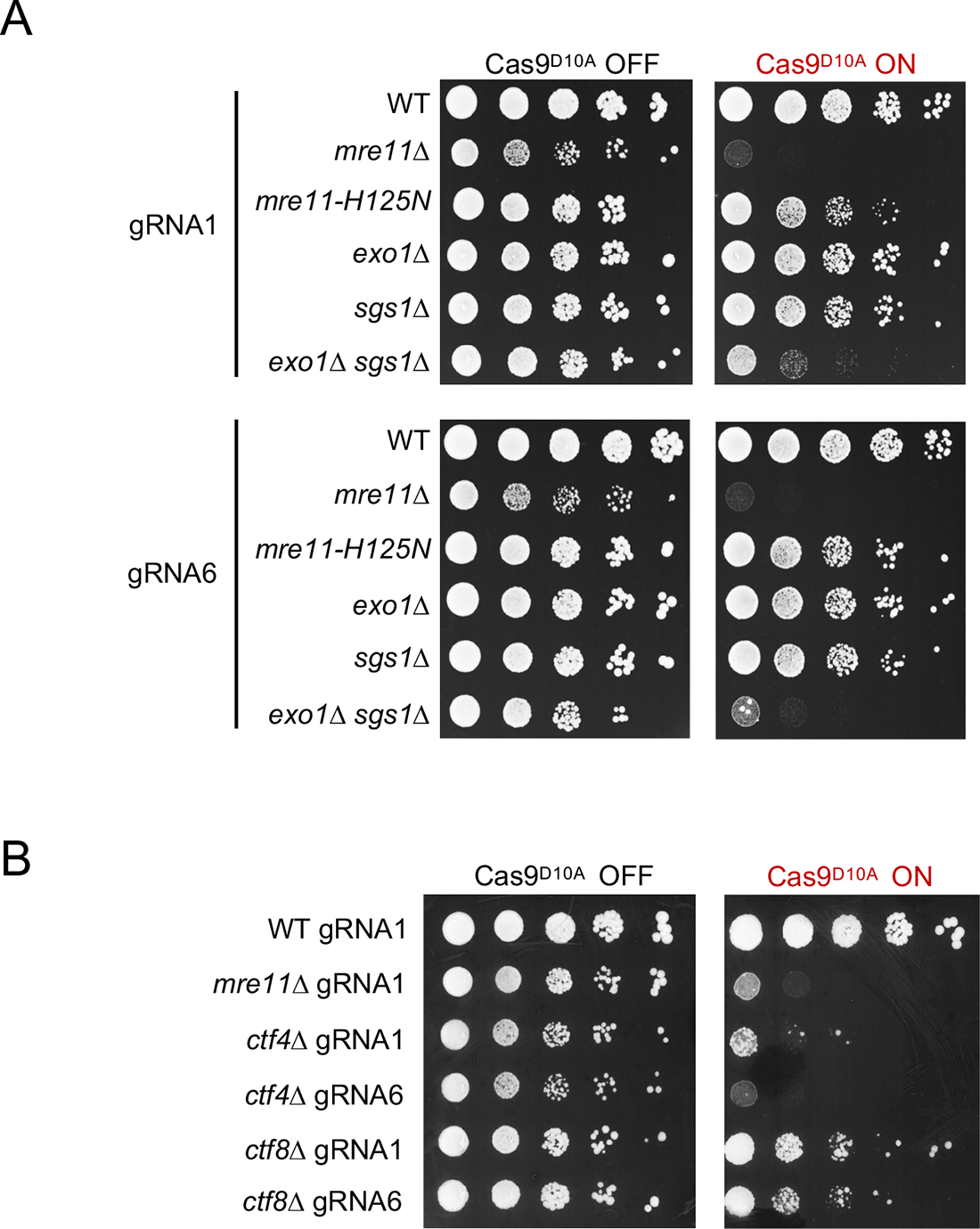
Long-range end resection and sister-chromatid cohesion are required for collapsed fork repair. A. Ten-fold serial dilutions of strains lacking Mre11 nuclease or long-range resection factors were spotted on YPD or YPD + β-estradiol. B. Ten-fold serial dilutions of strains lacking Ctf4 or Ctf8 were spotted on YPD or YPD + β-estradiol. All strains were transformed with plasmids expressing Cas9^D10A^ and gRNA1 or gRNA6, as indicated.

**Figure S4.**
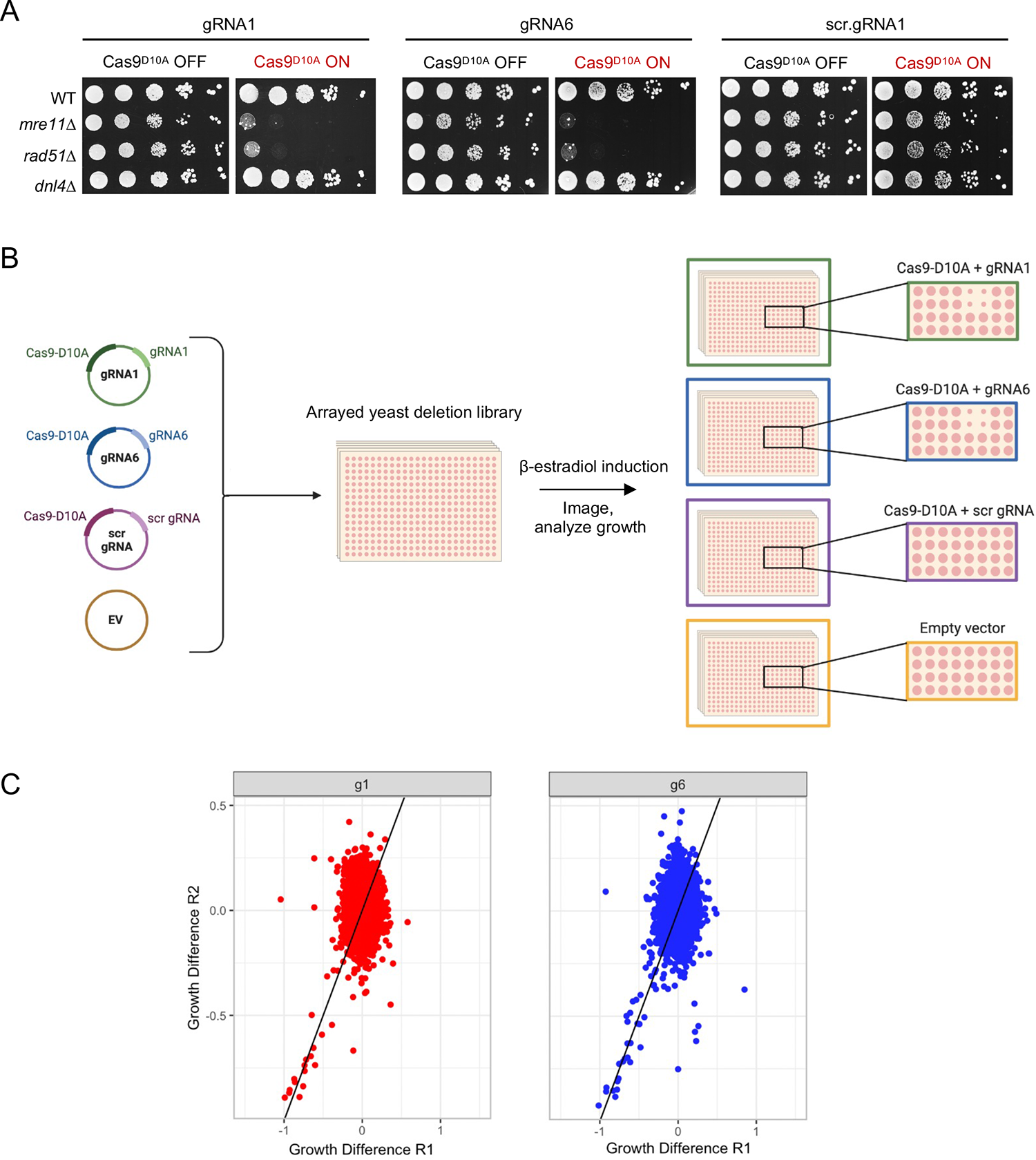
Strategy for genome wide screen in the S288C strain background. A Ten-fold serial dilutions of the WT, *mre11Δ*, *rad51Δ* and dnl4*Δ* S288C strains were spotted on YPD or YPD + β-estradiol. All strains were transformed with plasmids expressing nCas9 and gRNA1, gRNA6 or scr.gRNA1, as indicated. B. Schematic of the method used to introduce plasmids into the arrayed yeast deletion library and screening procedure. C. Biological replicates of the screen with gRNA1 and gRNA6. A line with a slope of 1 is plotted as a reference but does not represent correlation.

**Figure S5.**
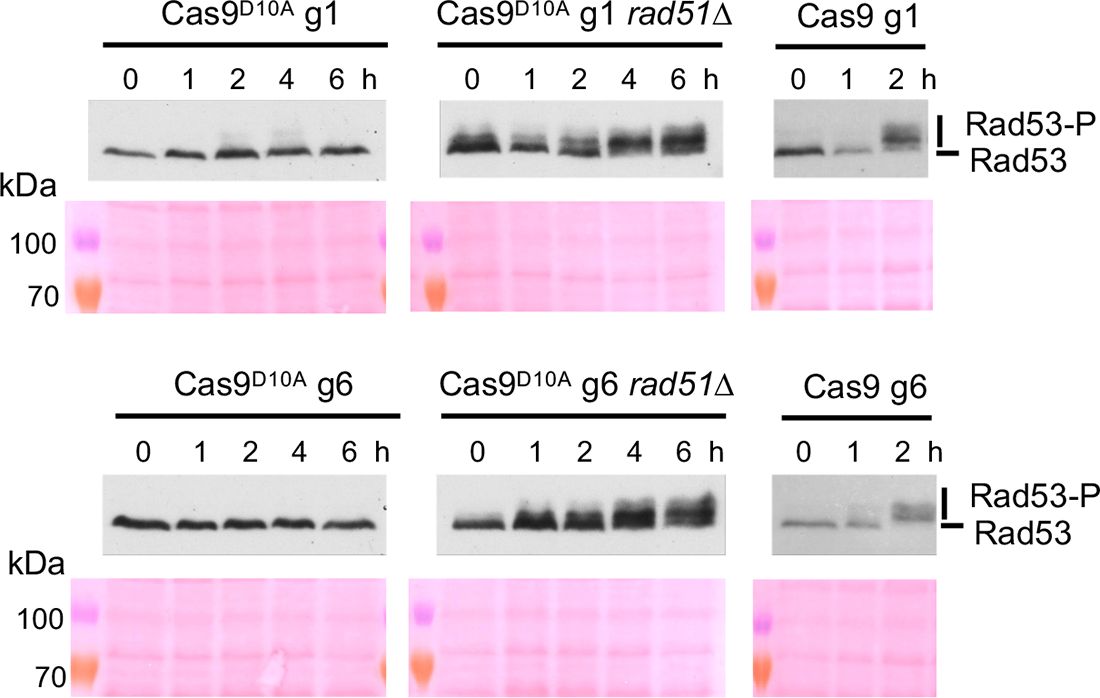
A single broken replication fork fails to activate the DNA damage checkpoint. Western blots to detect Rad53 phosphorylation (top) and corresponding Ponceau S staining (bottom).

**Figure S6.**
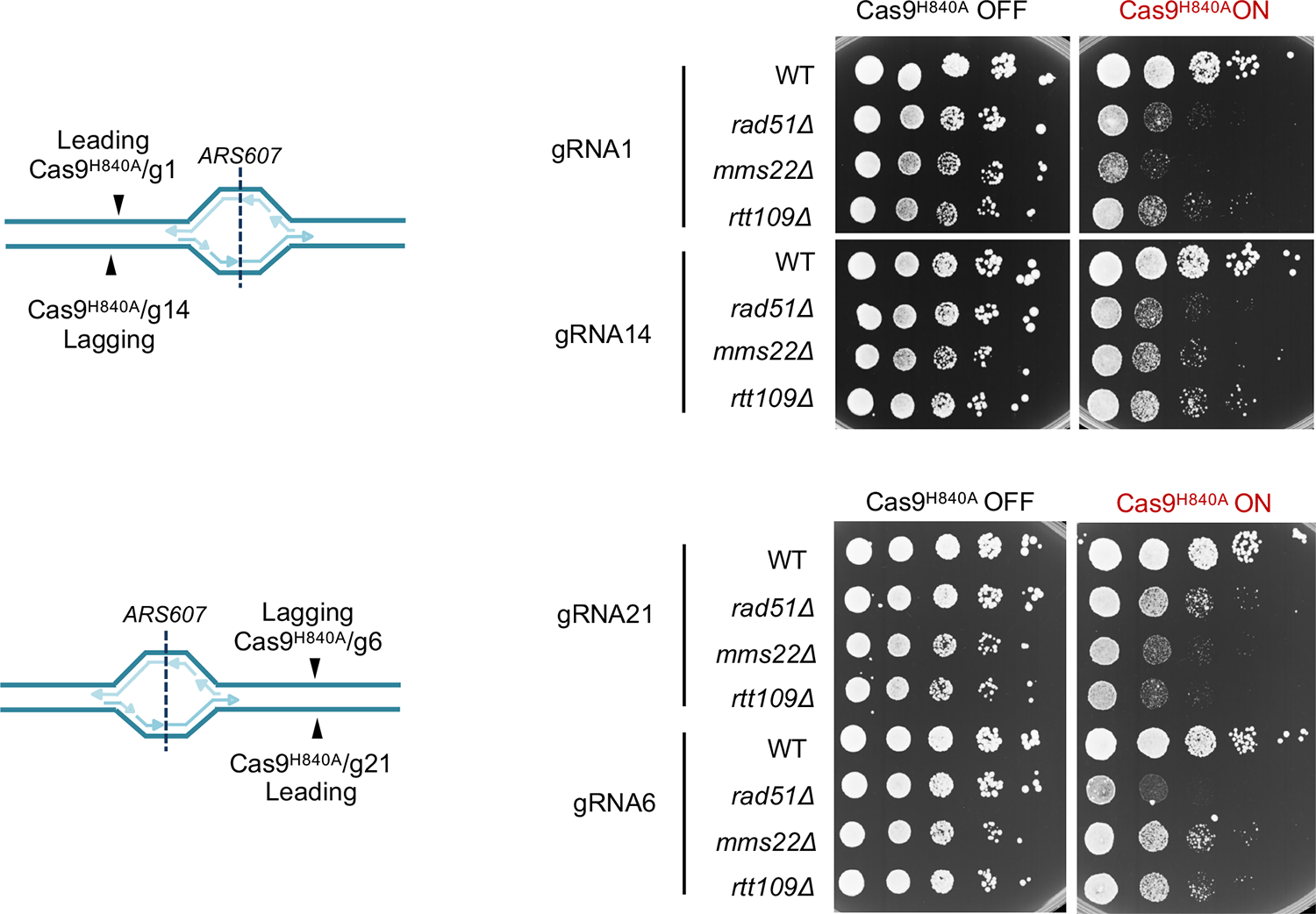
Strand specificity for RCNA function in collapsed fork repair. A schematic representation of nicked strand using the indicated nCas9/gRNA combination (left). Ten-fold serial dilutions of the indicated strains were spotted on YPD or YPD + β-estradiol (right).

**Supplementary Table 1.**
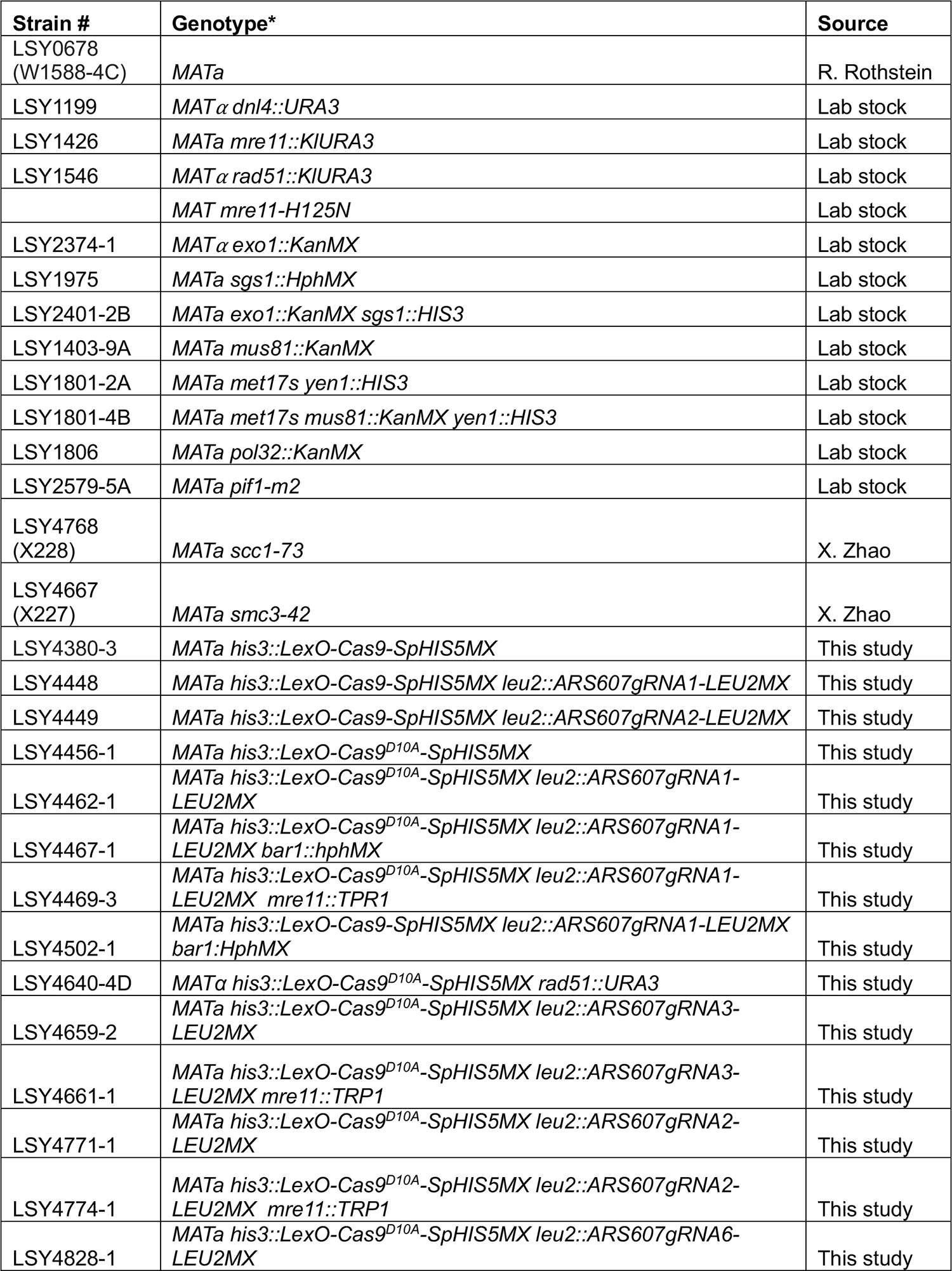
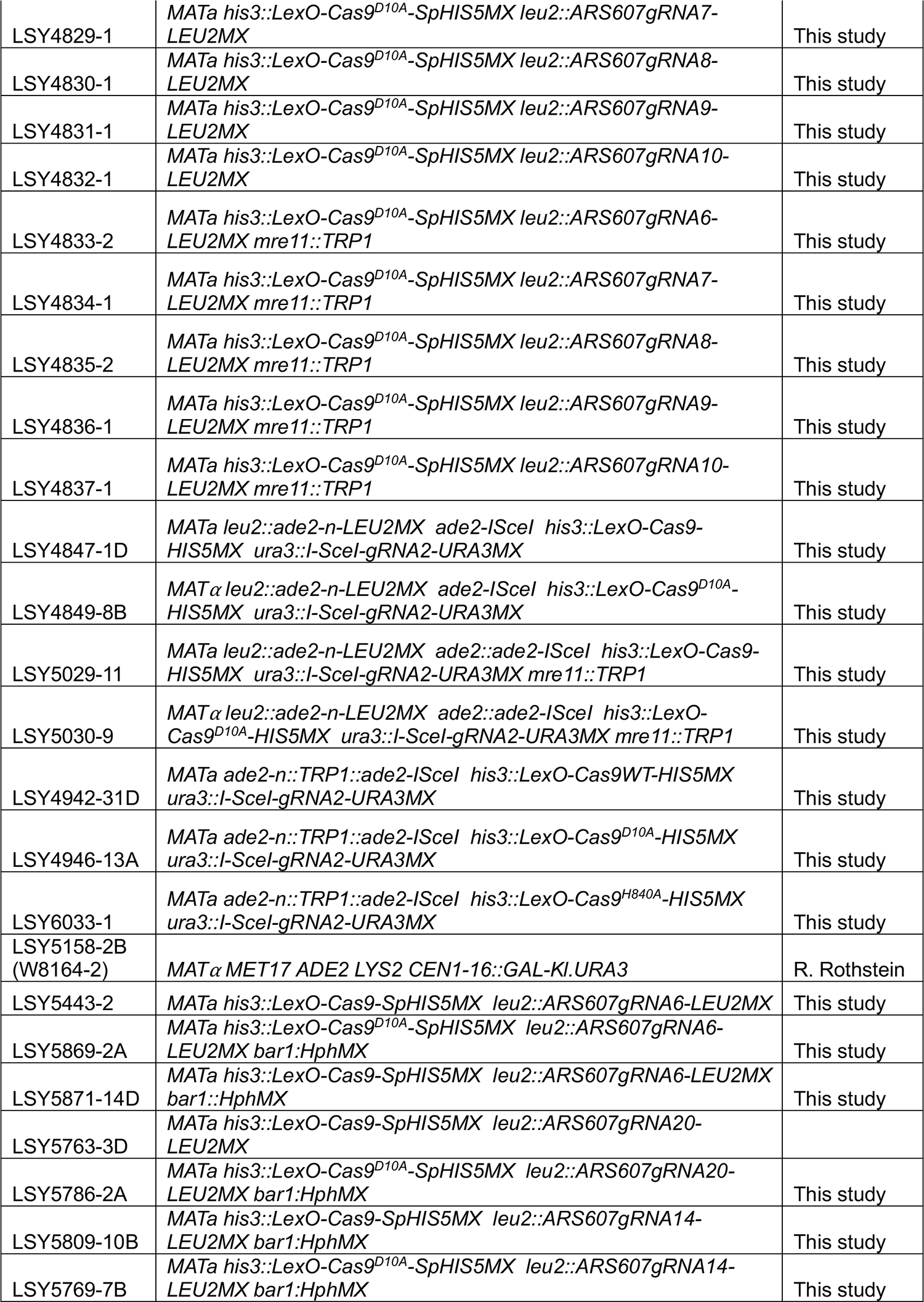
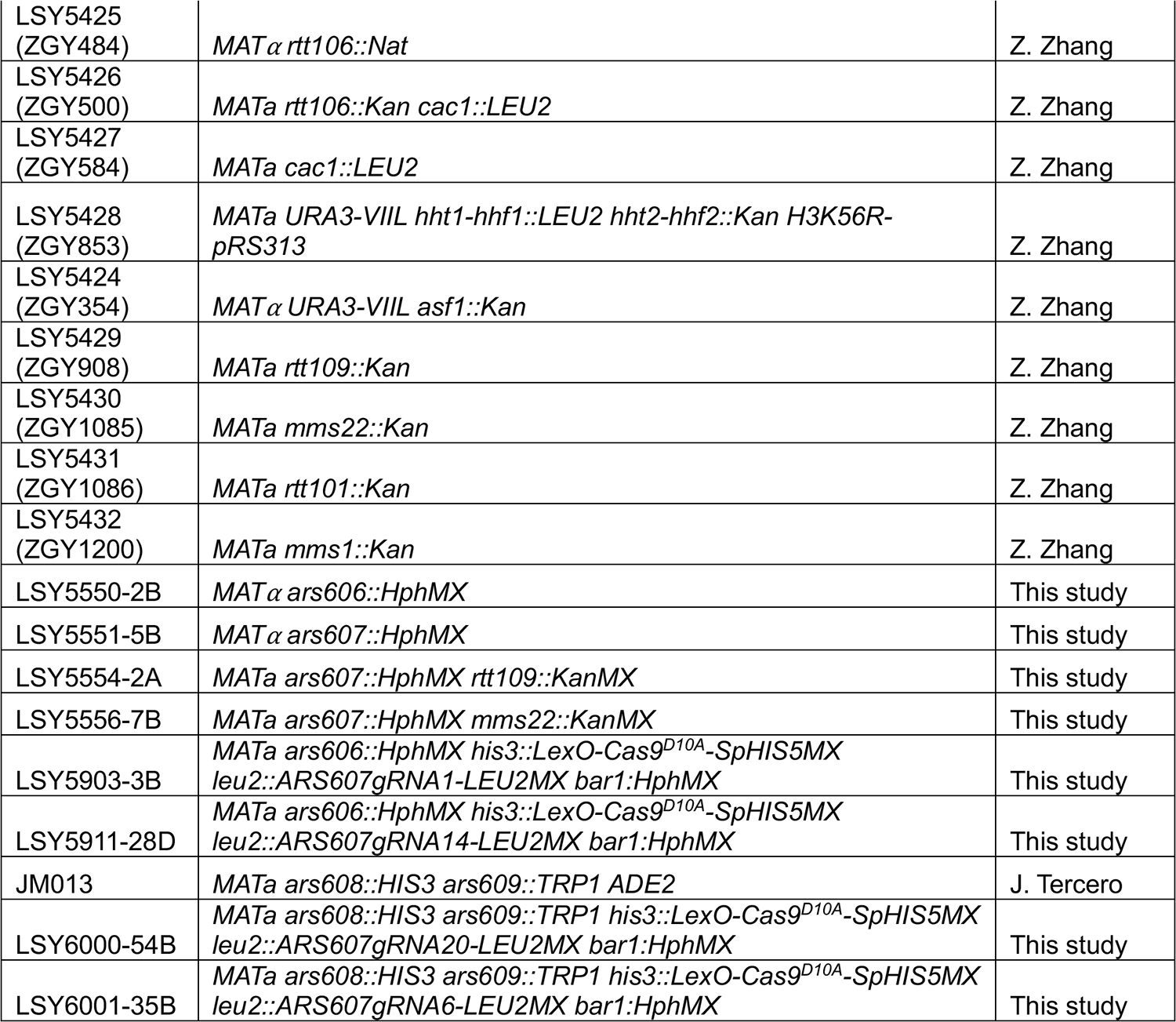
Yeast strains.

**Supplementary Table 2.**
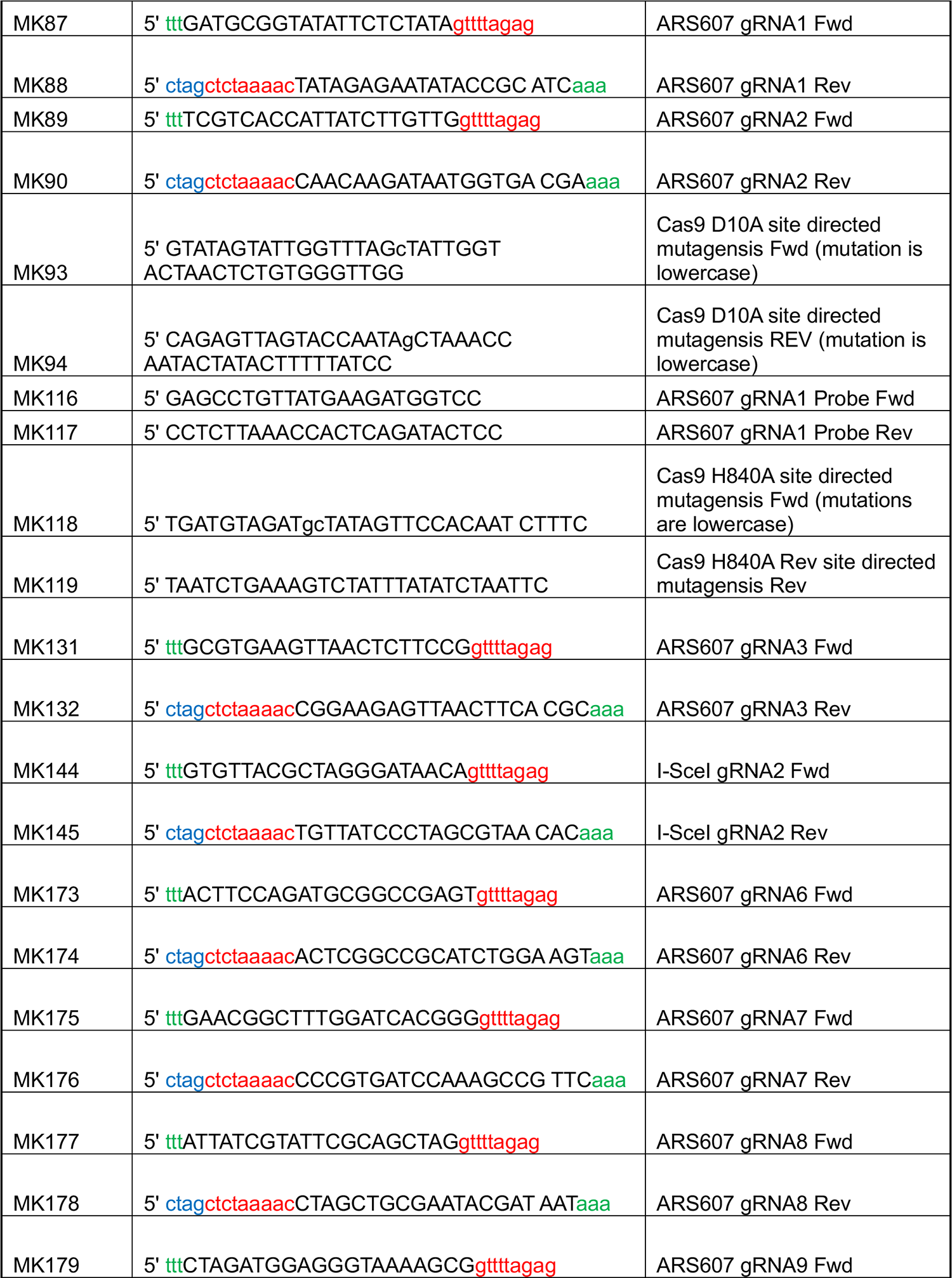
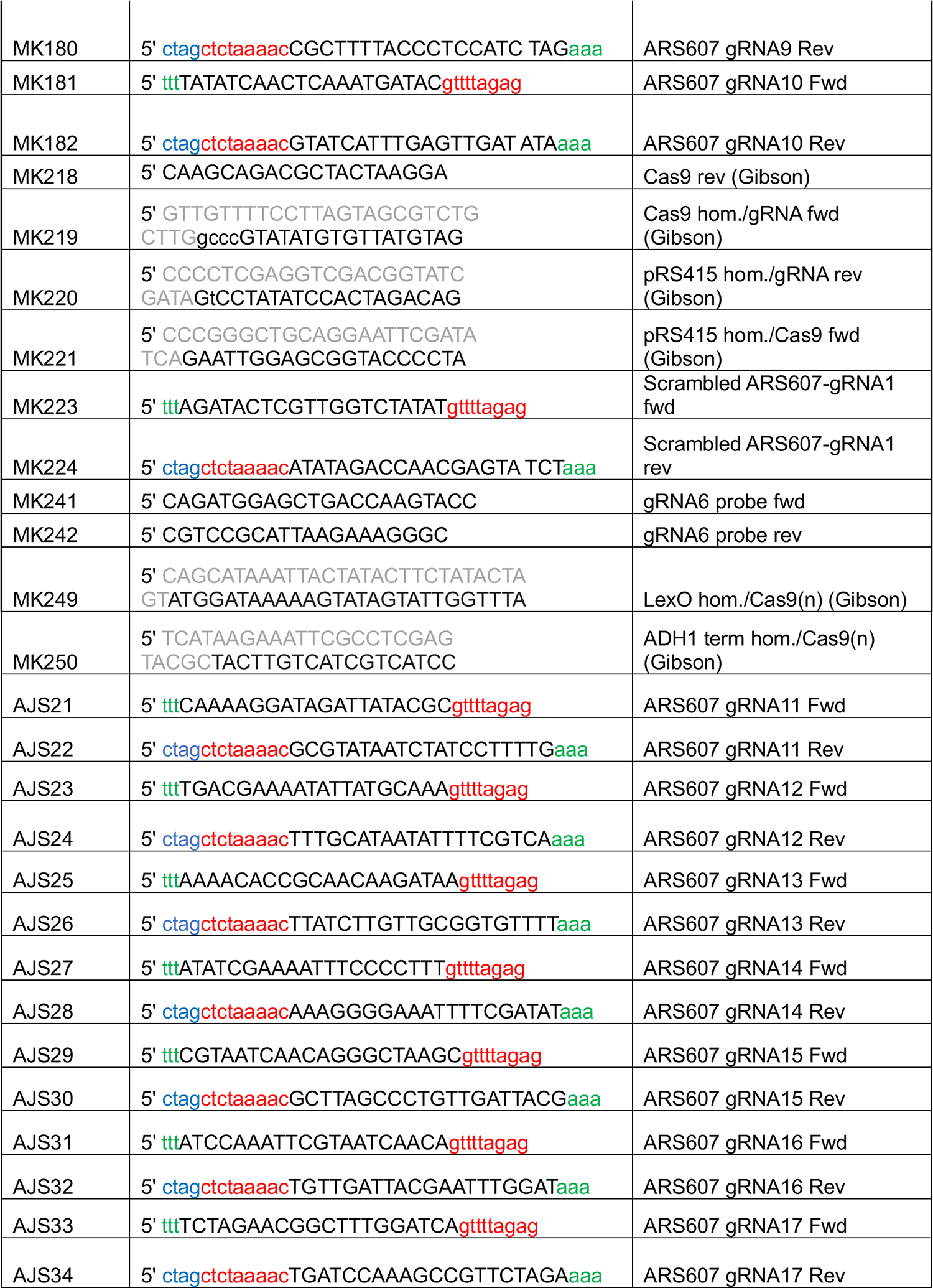
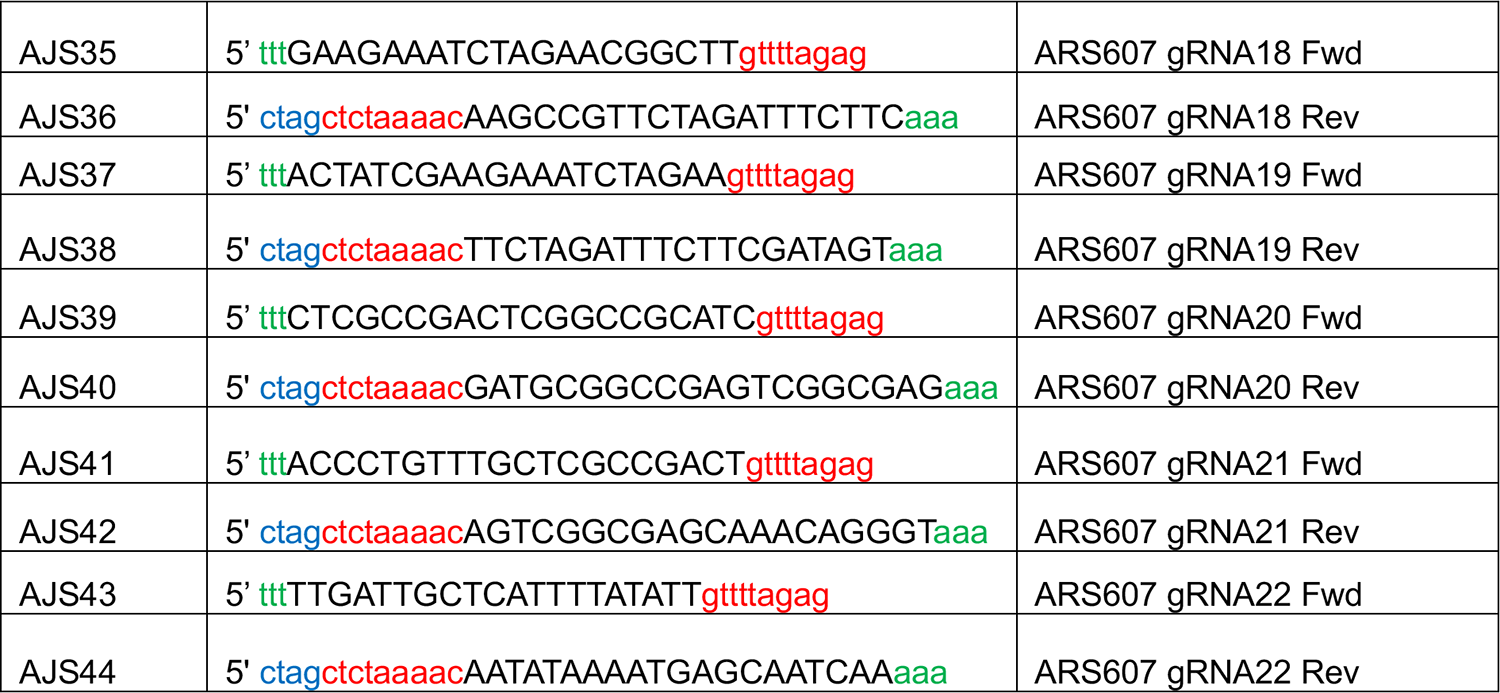
Oligonucleotides.

**Supplementary Table 3.**
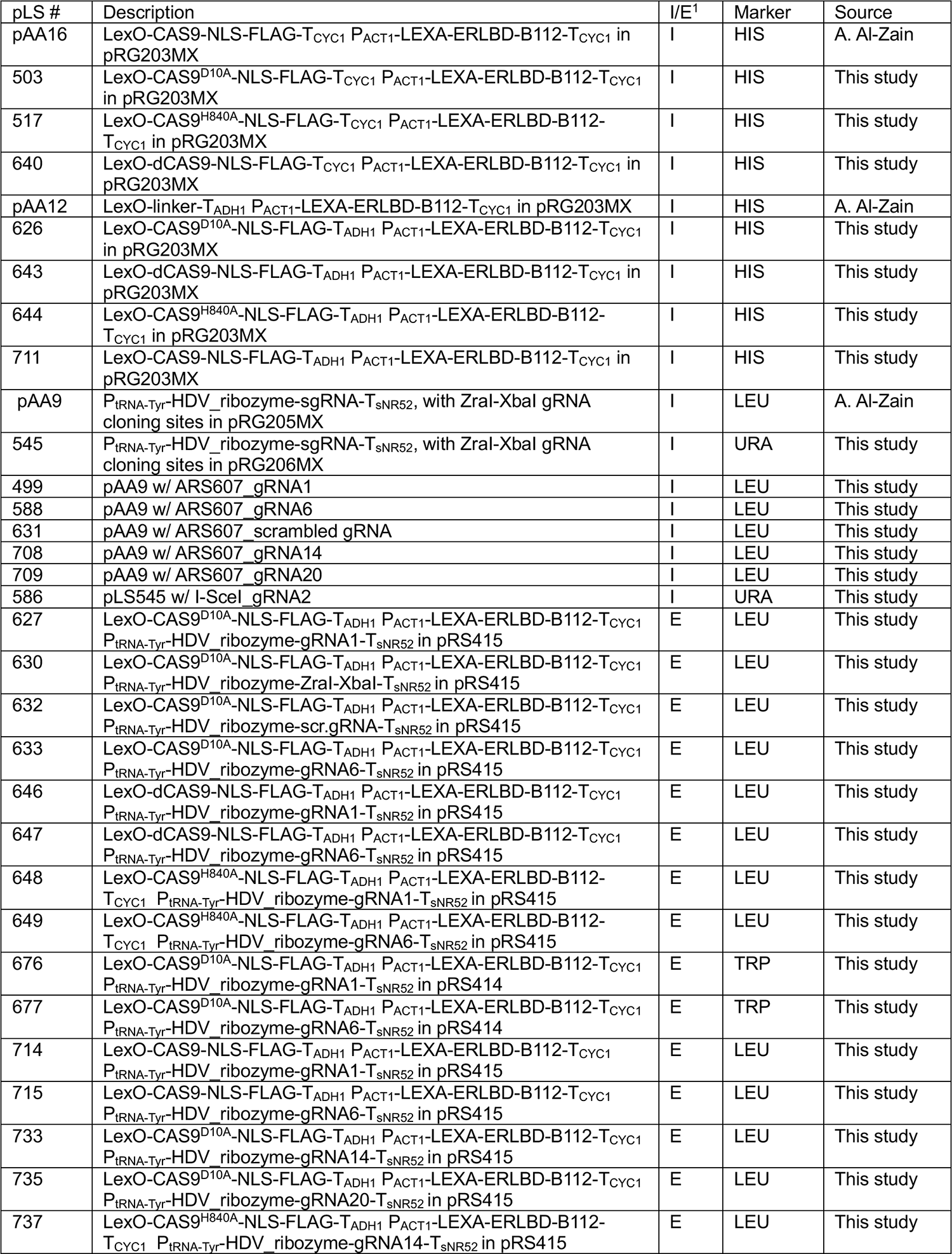
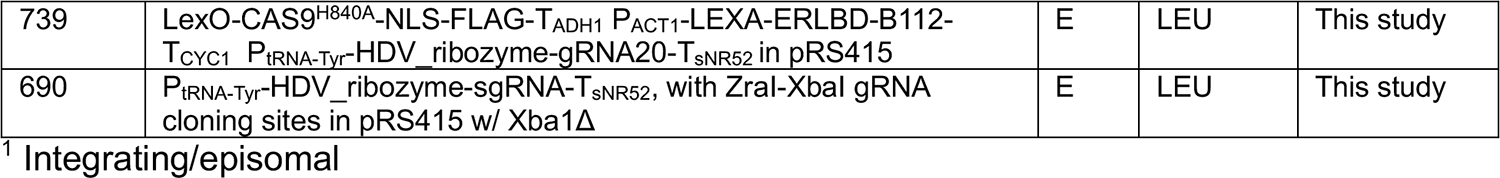
Plasmids.

